# A Gene-expression Module Identifies Circulating Immune Cells with Enhanced Recruitment to Sites of Inflammation

**DOI:** 10.1101/2023.04.11.536347

**Authors:** Debajyoti Sinha, Thomas Laurent, Alexis Broquet, Cynthia Fourgeux, Thibault Letellier, Gaelle Tilly, Sarah Bruneau, Simon Ville, Laurence Bouchet-Delbos, Julien Brancherau, Clarisse Kerleau, Sophie Brouard, Gilles Blancho, Magali Giral, Regis Josien, Richard Danger, Antoine Roquilly, Nicolas Degauque, Jeremie Poschmann

**Affiliations:** Nantes Université, CHU Nantes, INSERM, Center for Research in Transplantation and Translational Immunology, UMR 1064, ITUN; F-44000, Nantes, France; CHU Nantes, Nantes Université, Institut de Transplantation Urologie Néphrologie (ITUN), Service de Néphrologie et Immunologie Clinique, 44000 Nantes, France; Nantes Université, CHU Nantes, INSERM, Anesthesie Réanimation, CIC 1413, F-44000 Nantes, France

## Abstract

Circulating immune cells are critical mediators of inflammation upon recruitment to tissues, yet how their gene expression state influences this recruitment is not well understood. Here, we report longitudinal single-cell transcriptome profiling of peripheral blood mononuclear cells in patients undergoing kidney transplantation rejection. We identify a novel gene expression module, termed ALARM (early activation transcription factor module), associated with transcriptional regulation, homing, and immune activation across multiple immune cell types. Circulating cells expressing this module are significantly reduced in patients experiencing graft rejection, a finding confirmed in a pig model of acute kidney transplantation rejection. Correspondingly, module expression is markedly increased in kidney grafts undergoing rejection, indicating preferential recruitment of ALARM-expressing cells to the inflamed tissue.

Within this module, we identify the receptor CXCR4 and its ligand CXCL12, expressed in the graft, as a likely mechanism for recruitment. In vitro transwell assays combined with scRNA-seq reveal that this CXCR4-CXCL12 interaction is critical for T cell migration and upregulation of CD69, an early activation marker, and is accompanied by a metabolic switch towards glycolysis. Further exploration of publicly available transcriptomic data demonstrates that this module is generally expressed in healthy individuals and is strongly associated with responses to infection, including SARS-CoV-2 infection. This finding is further supported by experiments in a pneumonia mouse model, which confirm the recruitment of CXCR4-expressing T cells during lung infection. Moreover, we find that module expression is predictive of immune-mediated diseases.

In summary, we have identified a key gene expression module in circulating immune cells that orchestrates their preferential recruitment to inflamed tissues, metabolic reprogramming, promoting tissue residency and effector functions. These insights advance our understanding of immune cell recruitment and activation mechanisms in transplant rejection and infectious diseases, with potential implications for therapeutic interventions.

## Introduction

Circulating immune cells are critical to be recruited to the site of inflammation, infection, and cancer. This compartment and particularly the peripheral blood mononuclear cells (PBMCs) thus offer an attractive resource for precision medicine as in a single experiment, diverse cell types, including the CD4^+^ T cells, CD8^+^ T cells, B cells, NK cells and monocytes are profiled. Especially since the introduction of single cell RNA-seq (scRNA-seq), profiling PBMCs has been highly successful in identifying gene expression signatures and cell-types associated with immune-related diseases^1–4^. For example, a monocyte signature associated with sepsis was discovered in circulating cells ^1^. Recently, a study on systemic lupus erythematosus revealed gene expression changes with disease state and genetic variation ^2^. In addition, distinct Covid-19 studies revealed signatures associated to infection and disease severity in blood ^5,6^.

Transcriptomics profiling enables the characterization of genes groups (i.e., modules) which perform critical cellular functions such as maintaining a cell identity, homeostasis & metabolism and respond to external signals. Notably, in circulating immune cells we have previously shown that monocytes express a gene module associated with Herpes Simplex virus reactivation after traumatic brain injury^7^. While gene expression programs of circulating immune cells are likely to be distinct from the same cells which migrated into the tissue, identifying modules in circulating cells may reveal early immune activation programs or modules associated with homing and migration. For example, in a previous study we identified large gene regulatory and gene expression alterations in circulating monocytes during active Tuberculosis which improved pathogen clearance for these cells ^8^.

Leveraging on scRNA-seq and the availability of recent module identification approaches tailored for single cell transcriptomics ^9^ we aimed to identify gene expression programs associated with kidney transplantation rejection. Currently, rejection status is monitored in clinical practice by analyzing metabolites in blood and urine, such as creatinine, to assess renal function. The diagnosis is then confirmed through pathologic examination of kidney biopsies^10^. However, metabolite monitoring is not specific to rejection and can be approximative. Pathologic analysis of kidney biopsies remains the primary diagnostic tool, offering reliable results. However, despite minimal risk to patients, it is an invasive procedure that cannot be performed regularly. Thus identifying gene expression signatures in circulating cells may improve precision medicine diagnostics of kidney rejection ^11^. In addition, acute and chronic rejection are characterized by the infiltration of immune cells into the graft, which then mediate an inflammatory response in the tissue ultimately leading to the rejection of the graft. Blood thus constitutes an easily accessible compartment to identify gene expression modules associated with homing and early activation. Two archetypes of rejection are prominent according to the Banff classification^10^, the antibody mediated rejection (ABMR) and T-cell mediated rejection (TCMR), which can also arise in a mixed form. In both cases, immune cell infiltration into the graft occurs via the bloodstream through either donor specific antibodies binding to the graft endothelium in ABMR, or cytokine and homing signals in TCMR^12^.

In this study, our aim was to identify putative modules in circulating cells which may be associated to kidney transplantation rejection. We profiled a longitudinal patient cohort consisting of 3 stable patients, 3 TCMR and 3 ABMR patients at 0-month, 3 month and 12 months after transplantation or when rejection occurred. The PBMCs were collected at the same time of graft biopsy, allowing us to characterize relationship of graft rejection status with gene expression modules of peripheral immune cell-types. We identified a module associated with transcriptional regulation and early activation in the blood and used a pig-transplantation model to validate its association with rejection status. Further characterization of this module was carried out in transcriptomics data from over 1500 kidney biopsies revealing a cytokine-receptor interaction between the graft and circulating cells, respectively. Finally, we demonstrate that this module is not specific to graft rejection but is implicated and predictive of a variety of immuno-pathologies.

## Results

### Single cell transcriptome analysis of circulating immune cells in a longitudinal kidney transplantation cohort

In order to identify gene expression changes in circulating immune cells during kidney transplantation rejection, we used scRNA-seq on PBMC isolated from 3 patients with stable allograft function (STA) for which no sign of rejection was observed clinically after more than a year of follow-up, 3 antibody mediated rejection (ABMR) patients and 3 T-cell mediated rejection (TCMR) patients (Figure 1A, Table 1). The patients were selected based on their treatment, age, sex, and collection time. (See Table 1).

**Figure 1.**
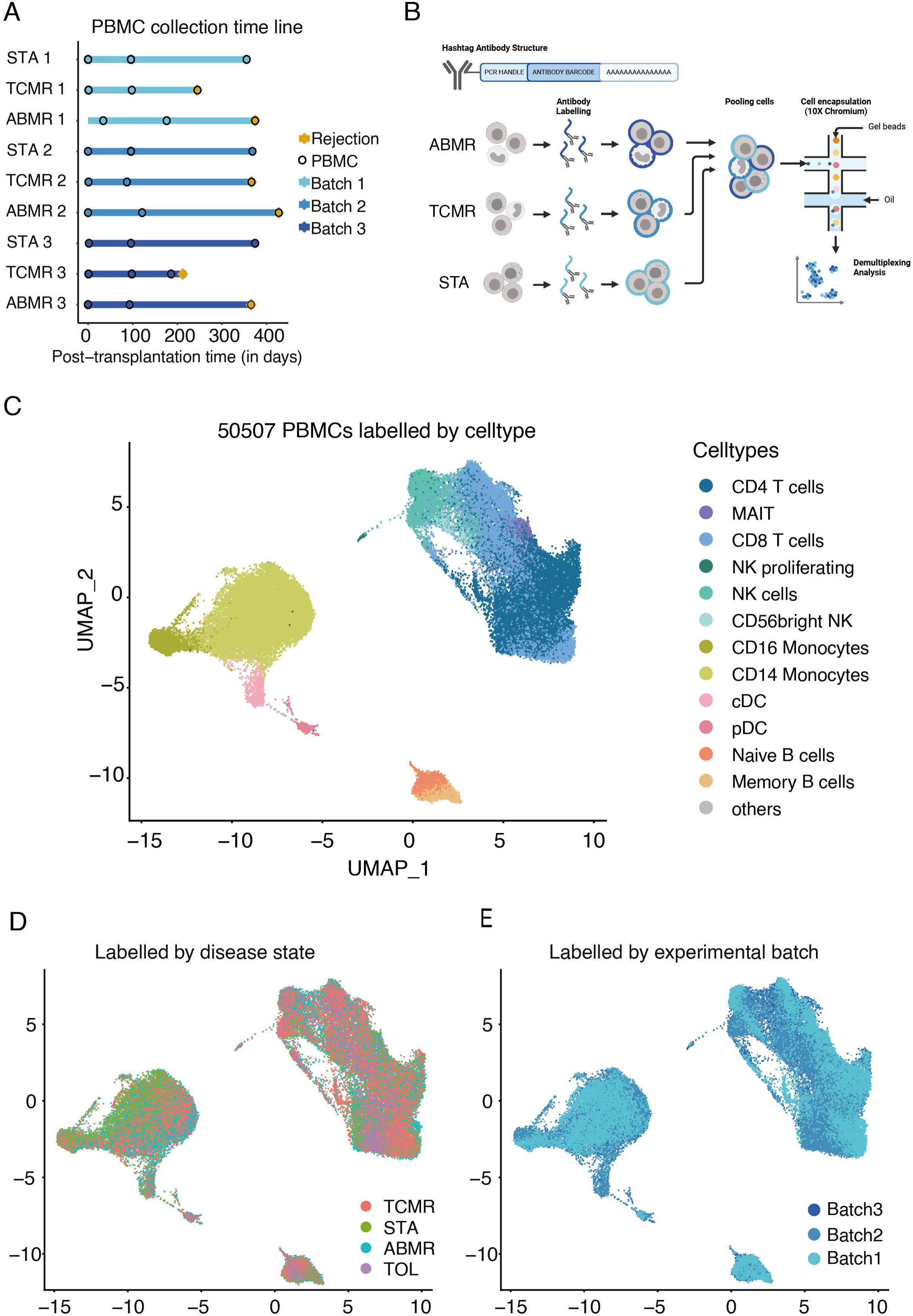
Comprehensive longitudinal single-cell RNA-sequencing of circulating immune cells in a cohort of kidney allograft recipient. A) Timeline of the blood sampling points post-transplantation for the patients followed longitudinally. STA=Stable patient (n=3), TCMR=T cell mediated rejection (n=3), ABMR=Antibody-mediated rejection (n=3). Tolerant patients are not shown. B) Schematic diagram of the scRNA-seq preparation workflow using cell hashing. Peripheral blood mononuclear cells (PBMCs) were collected from stable, ABMR and TCMR patients and then stained with one different oligo-conjugated antibody before being pooled and processed using microfluidic encapsulation. C) UMAP dimensional-reduction embedding of the integrated samples (n=30). Each colour represents a different cell subpopulation, adapted and manually curated from the automatic Azimuth annotation. D) UMAP projection showing the disease state distribution, TOL=Tolerant patients. E) UMAP projection coloured according to experimental batch of origin

**Table 1:**
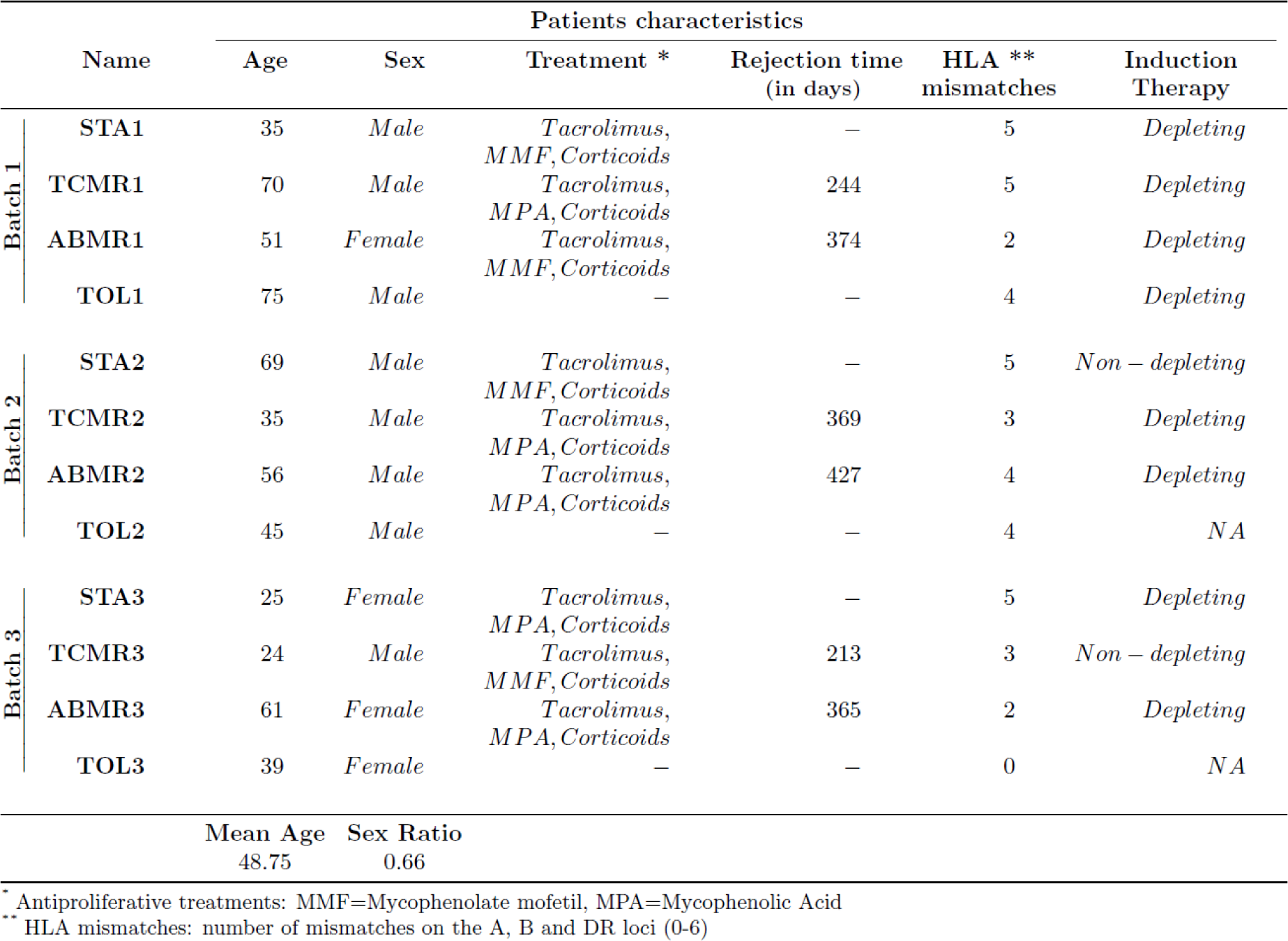
Clinical summary of the cohort composition.

For each patient, three time points were profiled; T0, at the kidney transplantation, T1 at 90-150 days after the transplantation and T2, which was sampled at the time of rejection for ABMR and TCMR or at 1 year for STA patients after transplantation (Figure 1A). In addition, PBMCs from 3 kidney transplantation patients who maintained allograft function in the absence of immunosuppression (i.e., operational tolerant (TOL)) were included in the cohort (Table 1).

Rejection status was defined by clinical pathology assessment of kidney biopsies performed at T1 and T2 for all patients. ABMR and TCMR were defined by pathology biopsy assessment at time T2 whereas patients were determined as STA when they which no sign of rejection in biopsies at T1 and T2. To minimize scRNA-seq-related experimental variation, we performed CITE-seq (Cell Hashing)^13^ using hashtag oligo-conjugated antibodies (HTO) to label each patient and time point separately and then pooled 10 samples (1 ABMR, 1 TCMR and 1 STA patient across T0, T1 & T2 and 1 TOL patient) into a single experiment. We thus generated the complete transcriptomic data in three balanced batches (Figure 1B). After removing doublets, data cleaning, normalization, and batch correction, we obtained 50,507 cells across the three batches (see Methods). Cells were automatically annotated using Azimuth^14,15^ and manually validated for cell-type specific markers (Figure 1C and Supplemental Figure S1A). Cell-type proportions varied minimally between conditions when compared to PBMCs from two separate cohorts of healthy volunteers (HV) from publicly available scRNA-seq data ^1,5^ (Supplemental Figure S1B). For example, NK cells were significantly decreased in the stable and rejection conditions as compared to HV. However, we note that there were also significant differences in the HV (e.g., CD14 and CD16 monocytes) from the two distinct control cohorts indicating that this may be due to individual variation. We then inspected the distribution of cells across patients and by time points (Supplemental Figure S1C) and across disease states (Figure 1D). Neither of these variables formed unique clusters suggesting that the clusters were driven by cell-type specific expression rather than by condition or cellular states as was also observed in other PBMC studies in patient cohorts ^1,2,5^. We then explored whether the clusters were affected by merging the three experimental batches (Figure 1E). As no batch effects were apparent through visual inspection, we used the KBET metric to quantitatively assess potential batch effects. KBET evaluates whether cells from different batches are clustering together in shared neighborhoods (i.e., clusters) (Supplemental Figure S1D)^16^. The acceptance rate of the KBET for the complete data set was 0.969, indicating that batch integration was successful. In summary, the pooling strategy and subsequent bioinformatics analysis resulted in a robust dataset of 50,507 cells to be analyzed for time and disease state specific gene expression.

### Gene co-expression analysis identifies a module associated to rejection state

To identify modules, ie. co-expressed groups of genes, we used gene co-expression analysis across all three batches independently (Figure 2A). This approach was chosen to avoid potential signal alterations induced during the batch correction step. We applied consensus non-negative matrix factorization (cNMF)^9^ to identify gene expression programs which may either be associated to cell-type specific gene expression programs or to cellular activity (see methods). Nine overlapping modules were identified which were evenly distributed across the three batches (Figure 2B, Supplemental Figure 2A). These modules mostly revealed cell-type specific expressions, notably three of these modules were associated to monocytes (Mod_1-Mod_3), and two modules were mostly expressed in a specific cell type such as B cells and pDC cells (Mod_4 and Mod_5 respectively) (Figure 2C). Three modules were enriched for the CD4 & CD8 T lymphocytes (Mod_6) and/or NK cells (Mod_7, Mod_8, Figure 2C). Interestingly, the Mod_9 was expressed in all cell types, but with notable higher expression in B cells, T cells, pDCs and NK cells as revealed by its module score (Figure 2C, Figure 2D).

**Figure 2.**
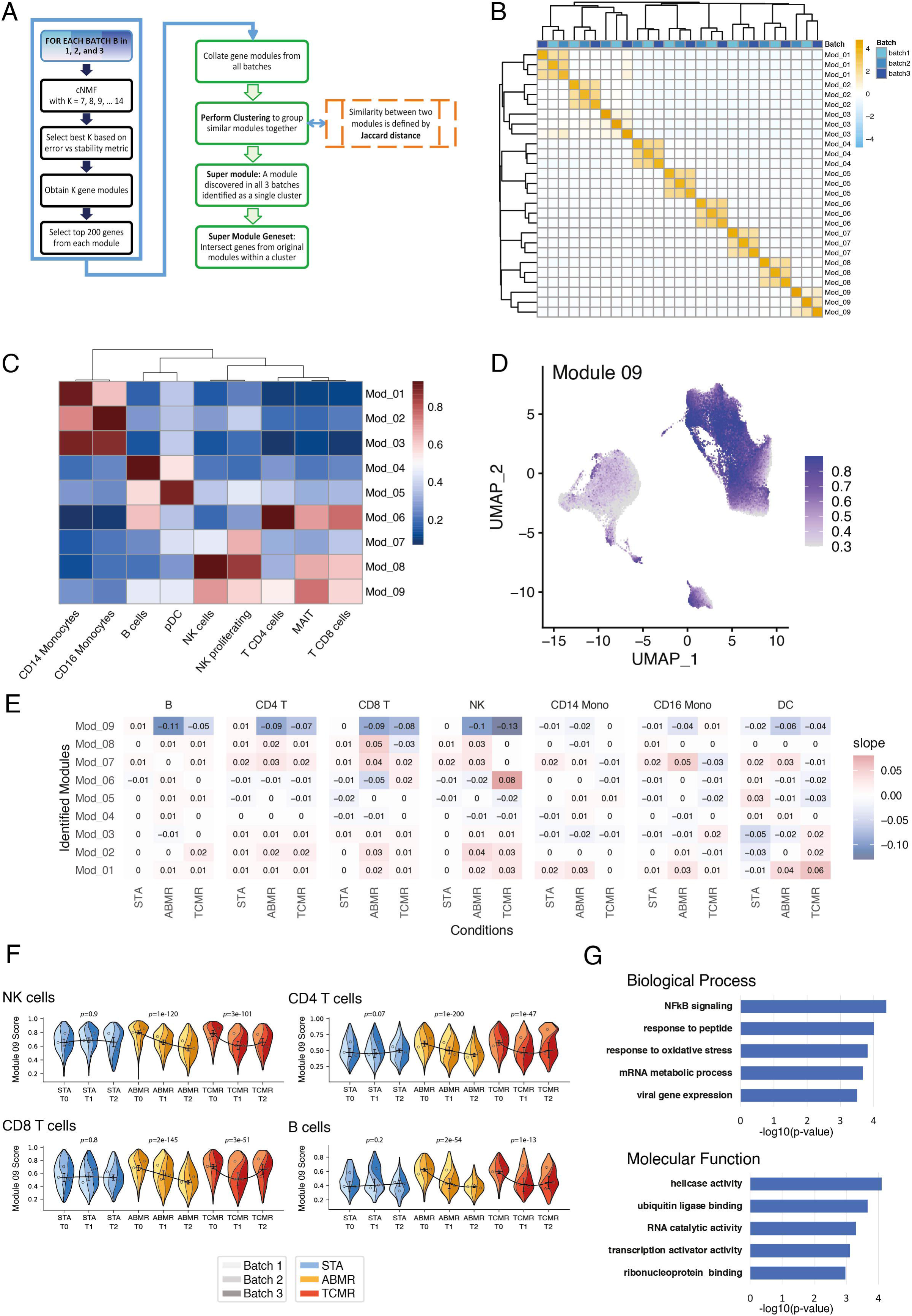
ALARM Module identification. A) Schematic workflow of the gene co-expression analysis using consensus nonnegative matrix factorisation (cNMF) module detection separate in each batch. The module selection was then refined based on overlapping genes between the three batches. B) Clustering of 9 modules across the 3 batches, using the Jaccard distance. C) Heatmap showing the combined gene expression of each module, i.e. module scores of the 9 modules summarized for each distinct cell-type. D) UMAP projection of the expression of Module 9 across all cells. E) Regression analysis of the 9 modules by cell types. The outcome variable (y) was time (T0, T1 and T2) and the independent variable (x) was the module score. Heatmap shows the beta values (trend) for each cell-type and disease state. Negative values correspond to a decrease of the module score across the three time points, positive values to an increase. F) Super violin plots^62^ showing the longitudinal trend of ALARM module in NK, CD4 T cells (top), CD8 T cells and B cells (bottom), stratified by individuals from each batch. G) Gene Ontology analysis of biological process (BP) and molecular function (MF) identified by comparing the ALARM gene enrichment using the 4000 most variable genes as background (See Supplemental file 1).

Next, we aimed to independently validate the co-expression of the 9 modules (Supplemental file 1). To achieve this, we quantified the gene co-expression using the pair-wise Pearson correlation coefficient R. As the module was identified using cNMF, Pearson correlation thus acts as an independent evaluation of gene co-expression. We note that this approach is inherent to weighted gene co-expression analysis (WGCNA), a prominent method to identify modules in bulk and single cell transcriptomics^17^. We compared the average Pearson correlation per cell for each module to the same number of randomly picked genes in the same cells. The eight cell type specific modules showed robust and significant correlation between module genes, and this correlation was strongest in the cell-types to which they were associated (Supplemental Figure S2B). The cell-activity module Mod_9 was also significantly correlated in all cell types (Supplemental Figure S2C). Of note, randomly chosen genes picked 1000 times revealed a Pearson correlation of 0 in all cell-types indicating that unrelated genes typically do not correlate with each other (Supplemental Figure S2B, S2C). Therefore, the modules identified above are robustly co-expressed, as inferred by the two most prominent module detection methods.

To test whether this cell-activity module or any of the cell-specific modules were associated to disease state, (i.e., rejection or stable) and if it would vary throughout time, we estimated the module score for each cell type and calculated the trend of the module score longitudinally in each cell-type (Figure 2E). A positive or negative slope thus indicates whether a module changes over time across the stable, humoral & cellular rejection. Indeed, Mod1 and Mod2 showed a significant positive trend in both rejection conditions (ABMR and TCMR) but not in the STA condition. Interestingly, the activity module (Mod_9) showed a significant negative trend (regression β value) in the rejection conditions but not in the stable condition in multiple cell types (B cells, CD4 and CD8 T cells and NK cells). Further inspection of this negative trend was carried out by displaying the module score of each patient separately in the form of a combined violin plot (Figure 2F). These module scores show that there was indeed a reduction in ABMR patients, while in TCMR patients the module score followed a U-shape, reduced more during T1 and increased again at T2. Interestingly, the 3 stable patient’s module scores remained consistent across time in NK, CD4 and CD8 and B cells. These results signify that Mod_9 expression is associated to rejection state in a time dependent manner.

### Discovery of the early activation, transcription factor module (ALARM)

To explore the function of the 61 genes found in Mod_9 we first investigated whether it was enriched for ribosomal, proliferation or cell cycle genes using SEURAT-based list ^18^. We did not observe any genes involved in these cellular processes (data not shown). We thus explored the genes within this module by performing gene ontology analysis. Enrichment of the module genes was quantified for molecular function (MF) and biological processes (BP) compared to the combined set of 4000 most variable genes from the three batches (Figure 2G). MF could be associated to 24 genes and was significantly enriched (FDR <0.05) for helicase activity, ubiquitin-like protein binding, RNA catalytic activity, transcription activator activity and ribonucleoprotein binding. The BP (20 genes) was associated to NF-κB signaling, response to peptide and oxidative stress, regulation of RNA metabolic processes and viral gene expression. Of the 61 genes in the module, 56 were annotated in the GSEA database and 30 of these genes were linked to gene ontology enrichment. This indicates that Mod_9 is likely to be involved in multiple molecular functions associated with transcription, mRNA process and ubiquitination. The BP suggested involvement in response to immune conditions (i.e., viral gene expression, NF-κB signaling, oxidative stress). This result was further supported by the 5-fold enrichment of Transcription factor genes in this module (OR: 4.9; Fisher-Test P-value 2.5e^-5^), such as the AP-1 complex (*JUN, JUND, FOS*), *REL* (NF-κB subunit), *MAFF* and *NR4A2* (see methods, Supplemental file 1). Further manual examination exposed the early activation marker CD69, a cell surface type II lectin. This receptor was described to be rapidly expressed at the membrane in T cells upon TCR activation^19^. In addition, CD69 has been described as a marker of tissue retention of T cells ^20–22^. Interestingly, *CD69* gene promoter is controlled by AP-1 TF complex and NF-κB, both of which are also members of this module^23,24^. We also found the cell surface marker CXCR4 in this module, a chemokine receptor known to play a role in recruiting CXCR4 positive cells to the kidney after an ischemic injury via the chemokine CXCL12^25^. The role of the CXCR4/CXCL12 axis in kidney rejection is still unclear^26^ but an elevated expression of CXCL12 has been described in chronic kidney rejection^27^ suggesting it may act as a chemotactic signal to recruit immune cells in the inflamed tissues. In summary, the Mod_9 module comprises genes implicated in the response to stress, mRNA processing, early activation, and tissue-homing. For clarity, we named this module ALARM, which stands for for eArLy activation trAnscription factoR Module.

### Circulating ALARM cells are recruited to the graft during acute graft rejection in a pig kidney transplantation model

We found that circulating cells expressing ALARM are altered in ABMR and TCMR in a timely fashion (Figure 2). To independently validate this observation, i.e., whether ALARM high expressing cells are depleted in the circulation during kidney transplantation rejection, we took advantage of an acute rejection pig kidney transplantation model (Figure 3A). We decided to use this pig kidney transplantation model as they share anatomical, physiological and genetic similarities to human and offer the advantage to have well defined swine leukocyte antigen (SLA) genotypes ^28^. Two SLA-mismatched pigs were subjected to kidney transplantation, keeping one of their own kidneys (see methods). This model typically results in an acute TCMR within a few days after transplantation, as no immunosuppressive treatment is given (Figure 3A). Kidney biopsies and PBMCs were collected daily before and after the transplantation. Microscopical analysis of the biopsies at D2, D4 and D6 indicated a time-dependent infiltration of immune cells, culminating at day 6 (Figure 3B). This infiltration was quantified in three areas (excluding glomeruli) of each biopsy time point (see methods, Supplemental Figure S3A). Cell counts drastically increased from day 2 to D6, indicating a continuous immune cell accumulation over time (Figure 3C). We noted that the second pig did not display any signs of rejection, possibly due to early arterial ischemia of the transplanted kidney and it was thus discarded from the subsequent analysis. PBMCs collected at D0, D2, D4 and D6, were pooled in a single scRNA-seq experiment (see methods), resulting in a total of 4,411 annotated cells across cell-types and time-points (Figure 3D and Supplemental Figure S3B). We found that cellular proportions within the PBMC compartment drastically changed from D0 to D2, characterized by a drastic increase in monocyte proportions concomitant with a reduction of B, CD4+ and CD8+ T cells (Figure 3E). This suggests that the lymphocytes are rapidly recruited to the kidney graft and accumulate there, as demonstrated by the cellular invasion observed in the biopsies at the same time (Figure 3C). To test whether the decline of blood lymphocytes is accompanied by a reduction of ALARM high expressing cells, we quantified ALARM expression across the time-points in CD4+, CD8+, B cells and NK cells (Figure 3F). Interestingly, as soon as D2 the levels of ALARM expression drastically decreased and remained low in the blood until sacrifice of the animal (D6). Taken together, this controlled time-course experiment reveals a drastic immune cell infiltration in the graft associated with the depletion of ALARM high expressing cells in the circulation. This result mirrors the reduction of circulating ALARM cells observed during the kidney transplantation rejection in the human cohort (Figure 2). It is thus probable that ALARM high expressing cells are readily circulating in healthy condition. Upon the kidney graft transplantation ALARM high expressing cells are then preferentially recruited to the organ to mediate the immune response.

**Figure 3.**
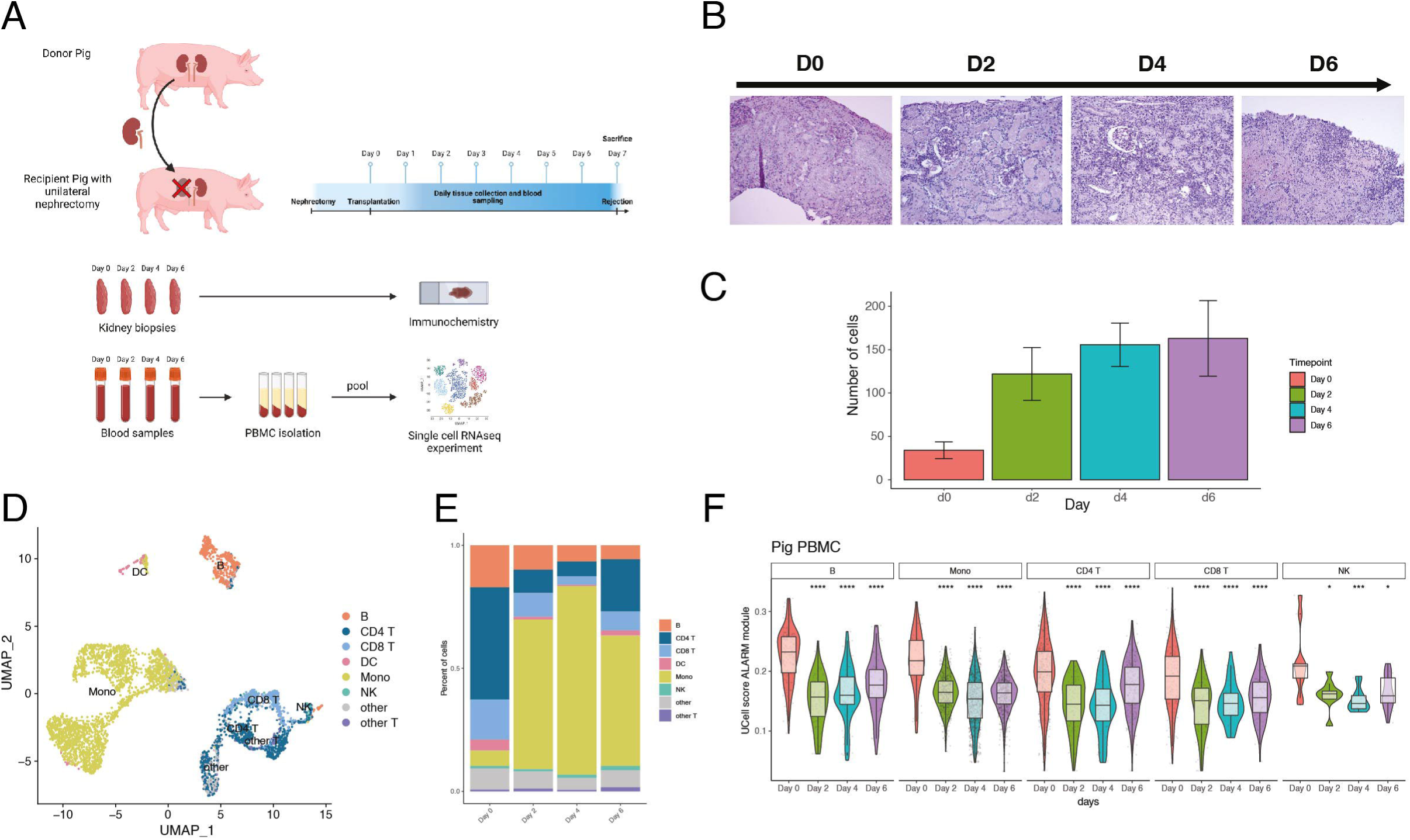
Pig model scRNA-seq analysis. A) Schematic diagram of the acute kidney allograft rejection in pig model. Recipient pig with a unilateral nephrectomy received a kidney graft from a second healthy pig. Kidney biopsies and PBMCs were collected daily and observed in immunohistochemistry. PBMCs were prepared for a scRNA-seq analysis. B) Immunohistochemistry of pig biopsies stained with Periodic Aid Schiff (PAS), to stain polysaccharides, glycoproteins and glycolipids at day 0, day 2, day 4 and day 6 C) Quantification of cell populations in the kidney graft at given days using Image J cell counting software (See supplemental figure 3 for details) D) UMAP dimensional-reduction projection of the circulating immune cell types (PBMCs) scRNA-seq after PBMC isolation at D0, D2, D4 and D6. E) Proportions of circulating immune cells (PBMC) across the different time points (D0-D6) in the recipient transplanted pig. F) Violin plot of the ALARM module score in B cells, monocytes, CD4 T cells, CD8 T cells and NK cells throughout acute rejection in the kidney tissues (Day 0, Day2, Day 4 Day6). \**P* < 0.05, \*\**P* < 0.01, \*\*\**P* < 0.001, \*\*\*\**P* < 0.0001.

### ALARM gene expression increases in kidney grafts undergoing rejection

To further support the hypothesis that ALARM high-expressing cells are recruited to the graft during rejection from the bloodstream, we evaluated the expression of ALARM genes in graft biopsies from kidney transplantation patients. For this, we used a previously published Canadian transcriptomics analysis of 569 transplant biopsies collected from 13 clinical sites and with a patient classification of STA, TCMR, ABMR and mixed rejection (TCMR and ABMR)^29^. In parallel, we exploited a second similar Belgian transcriptomics study performed on kidney biopsies in 224 patients who were either stable (168 patients) or undergoing ABMR^30^. After precleaning and QC controls of the available microarray data (Methods), we quantified the ALARM gene expression in each cohort separately (Figure 4A). The ALARM score was consistently and significantly elevated in all three rejection cases compared to stable biopsies. Similarly, in the second study, ABMR samples showed a significant increase in ALARM expression, regardless of the presence of donor-specific antibodies. Analysis of the ALARM genes ranked by expression, further revealed that this score is driven by the upregulation of a large fraction of the ALARM genes, including *CD69*, *CXCR4*, *JUN* and *IRF1* in both cohorts (Figure 4B). Quantification of the ALARM module across rejection and stable revealed a significant upregulation of ALARM expression in both data sets (Figure 4C). These results indicate that ALARM genes are significantly increased during graft rejection across over 793 biopsies in two distinct studies. Given that graft rejection is defined by immune cell infiltration and that circulating ALARM high cells are depleted at the same time, it is possible that ALARM expressing cells are preferentially recruited to the graft to mediate the rejection. To investigate how circulating cells could be preferentially recruited to the graft, we explored the cytokine expression in the graft and receptor expression in the circulating cell subsets. We first identified all possible ligand-receptor pairs and then tested whether these pairs were differentially expressed between rejection and stable status. We found 10 differentially expressed cytokines in the graft pairing with 7 receptors upregulated in circulating immune cells in both cohorts independently (Figure 4D). The most prominent receptor was CXCR4, expressed in CD4, NK, CD8 and B cells. As mentioned above, CXCR4 is a member of ALARM genes, indicating a likely mechanism of signaling from the graft via CXCL12 and recruitment of ALARM cells expressing *CXCR4*. This cytokine receptor pair has been previously described as a homing mechanism in various distinct tissues, such as bone marrow^31,32^ and in cancer^33^. To test the relationship of this interaction, we quantified *CXCL12* expression in the graft biopsies stratified by ALARM expression (low (<25%), mid (25-75%), high (>75%)) and found that there was a significant increase in CXCL12 expression in the ALARM high group in both cohorts (Figure 4E). In summary, these results show that ALARM is increased during rejection in the graft, which strengthens the notion that circulating cells with high ALARM expression are preferentially recruited to the kidney graft via the homing signaling axis of CXCL12 and CXCR4.

**Figure 4.**
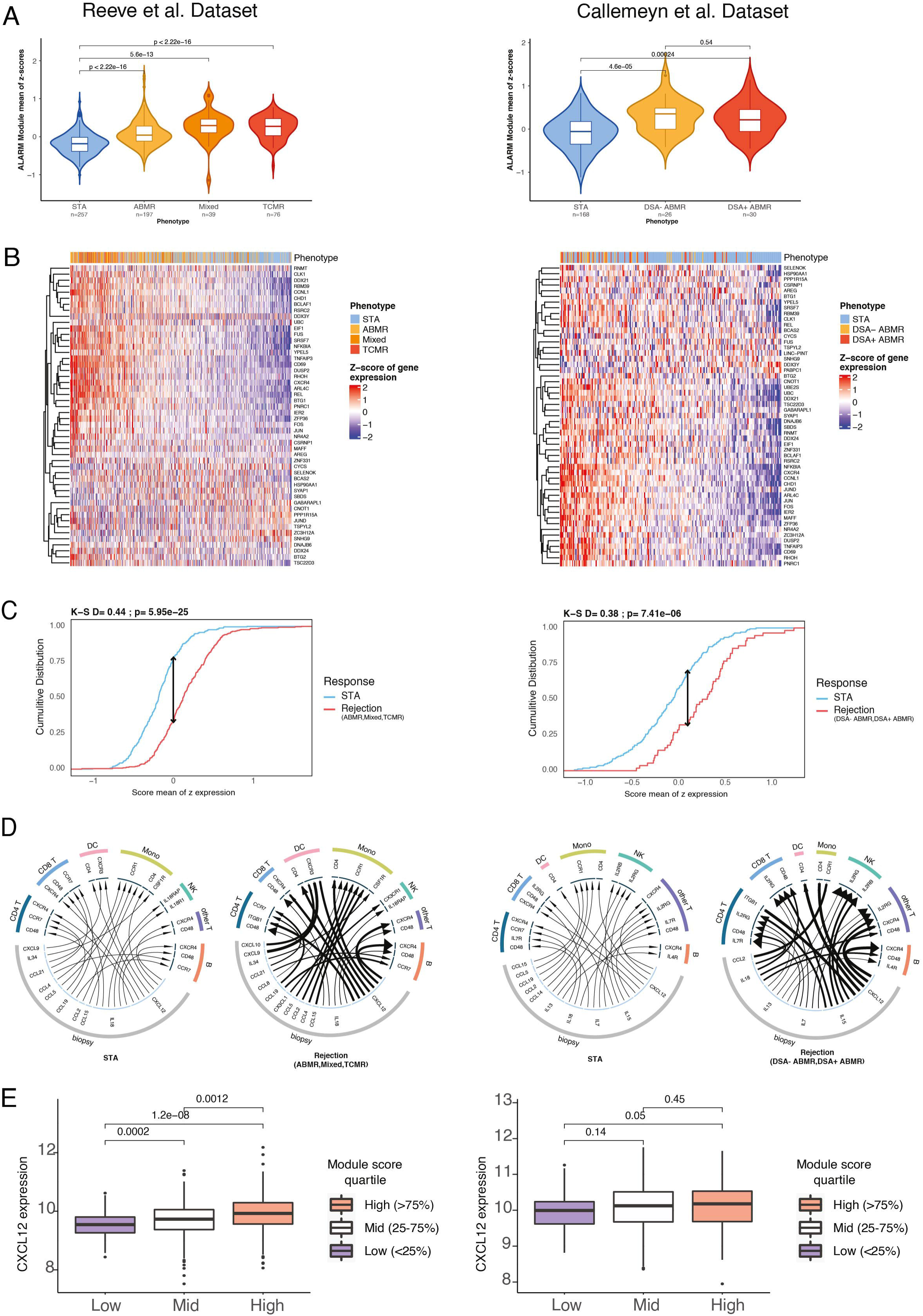
ALARM gene expression increases in kidney tissues during rejection. Left panels from Reeve et al., 2017 and right panels from Callemeyn et al 2020. STA =Stable patient, ABMR=Antibody-mediated rejection, TCMR=T cell mediated rejection, Mixed=Graft undergoing ABMR and TCMR, DSA-= Donor specific antibody negative and DSA+= Donor specific antibodies positive. A) Violin plot of the sum of z-scores of ALARM genes across conditions. Wilcoxon P-values are shown in panel comparing Stable (STA) to rejection status (ABMR, Mixed or TCMR and DSA- and DSA+ ABMR). The mean comparison p-values were computed using the Wilcoxon rank-sum test. B) Heatmap showing z-scores of ALARM genes (one gene per row) in all graft biopsies. Phenotype denotes the disease states. Patients are sorted on the mean of the module gene z-scores. B) Cumulative distribution of the mean of z-scores of ALARM genes comparing stable vs the combined rejection conditions. K-S = Kolmogorov - Smirnov P-values and distance. C) Ligand-Receptor analysis between receptors genes identified in circulating immune cells and cytokines genes expressed in the allograft kidney tissue under no rejection and rejection condition. Width of the arrow line is proportional to the expression of the ligands and the width of the arrowhead is proportional to the receptor. Only the top 5 associations from each cell type of differentially expressed receptors (PBMC scRNA-seq) and cytokines (Biopsy microarray) are shown. D) Boxplots showing CXCL12 expression in biopsies from patients with high ALARM module expression (>75%), medium (25-75%) and low (<25%). Wilcoxon rank-sum test was used to calculate P-values shown above.

### Analysis of ALARM Module Expression in an In Vitro Transwell Assay

To experimentally verify the observed CXCL12-mediated recruitment, we employed an *in vitro* transwell assay using a cytokine gradient of CXCL12. This assay consists of a membrane covered with primary microvascular endothelial cells (HDMEC), allowing for the comparison of unstimulated cells, cells in direct contact with CXCL12, and those migrating based on the CXCL12 gradient (Figure 5A).

**Figure 5:**
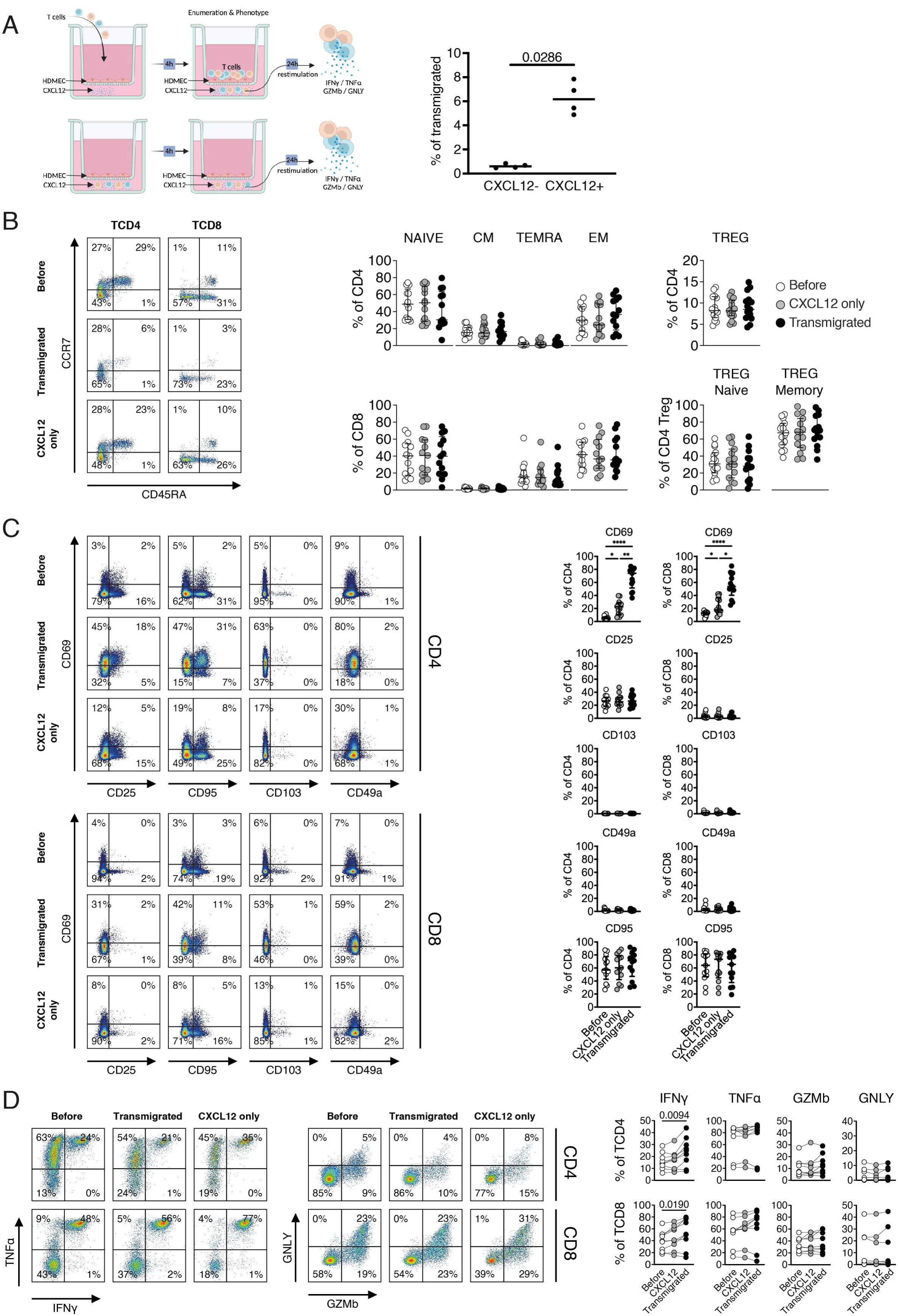
Analysis of ALARM Module Expression and T Cell Behavior in an In Vitro Transwell Assay. A) Schematic representation of the in vitro transwell assay used to study CXCL12-mediated recruitment of T cells. The assay involved a membrane covered with human dermal microvascular endothelial cells (HDMEC), allowing for the comparison between unstimulated cells, cells in direct contact with CXCL12, and those migrating across the membrane in response to a CXCL12 gradient. B) Quantification of T cell subsets following migration. The composition of naïve, central memory (CM), effector memory RA (TEMRA), and effector memory (EM) T cells remained similar post-migration and in response to CXCL12 alone, indicating that all T cell subsets are attracted to and migrate in response to the CXCL12 gradient. C) Analysis of T cell surface marker expression. CD69, a component of the ALARM module, showed a slight increase in response to CXCL12, with a more significant upregulation in migrated cells. CD25, CD49A, and CD95 levels remained unchanged. D) Functional analysis of migrated T cells. CXCL12-exposed and transmigrated cells were stimulated with PMA/ionomycin after 24 hours. Migrated cells showed significantly increased expression of IFN-γ in both CD4+ and CD8+ T cells, while TNF-α, granzyme B (GZMB), and granulysin (GNLY) expression remained constant Statistical significance was determined using one-way ANOVA, followed by post-hoc tests where appropriate. Significance levels are indicated as follows: *< 0.05, ** < 0.01, and *** < 0.001.

Flow cytometry quantifications of T cells indicate that CXCL12 significantly induces the recruitment and migration of T cells, observed 4 hours post-deposition (Figure 5A; p=0.0286). The composition of naïve, central memory (CM), effector memory expressing CD45RA (TEMRA), and effector memory (EM) T cells remained similar after migration and in response to CXCL12 alone (Figure 5B, Supplemental Figure S4A). This suggests that all T cell subsets are equally attracted and migrate in response to a CXCL12 gradient. We further analyzed the expression of several T cell surface markers, focusing on CD69 as a component of the ALARM module, CD25 (an activation marker), CD49a (a tissue residency marker), and CD95 (an apoptosis marker) (Figure 5C). Interestingly, CD69 increased slightly with CXCL12 addition but was even more highly upregulated in migrated cells, while CD25 expression did not change, decoupling the role of CD69 in early activation from its role in tissue residency^34^. CD49a and CD95 levels remained unchanged, indicating that CXCL12 in combination with migration specifically induced the extracellular display of CD69. It is noteworthy that CD69 expression depends on both CXCL12 and the direct contact with HDMEC cells (Supplemental Figure S4B).

To assess whether CXCL12-induced migration altered the T cells’ response to immune stimuli, CXCL12 exposed or transmigrated cells were purified and restimulated polyclonally for an additional 24 hours (Figure 5D). Migrated cells showed a significantly increased expression of IFN-γ in both CD4+ and CD8+ T cells in contrast to TNF-α, granzyme B (GZMB) or granulysin (GNLY). The increased expression of IFN-γ suggests that migration via the CXCL12 gradient may enhance the effector functions of T cells but that other signals are needed to induce cytotoxic mechanisms in this model. The observed increase in CD69 membrane display indicates a functional role for the ALARM module, which enables T cells to acquire “new functions” in the tissue i.e. to establish residency via CD69 expression and increased IFN-γ expression.

To further investigate the impact of CXCL12 signaling on T cell migration and the role of ALARM module expression, we performed scRNA-seq using the transwell assay under three distinct conditions (Figure 6A). In the first condition (CXCL12-), T cells were placed below the transwell membrane without any exposure to CXCL12. In the second condition (CXCL12+), T cells were placed below the transwell membrane in direct contact with CXCL12. The third condition, (transmigrated), involved placing T cells above the membrane, which were then collected from below the membrane after migrating in the presence of CXCL12. This setup allowed us to assess the transcriptional changes associated with T cell migration in response to CXCL12 (Figure 6A). Notably, all T cell subtypes identified by gene expression were found in the three conditions (Supplemental Figure S4C).

**Figure 6:**
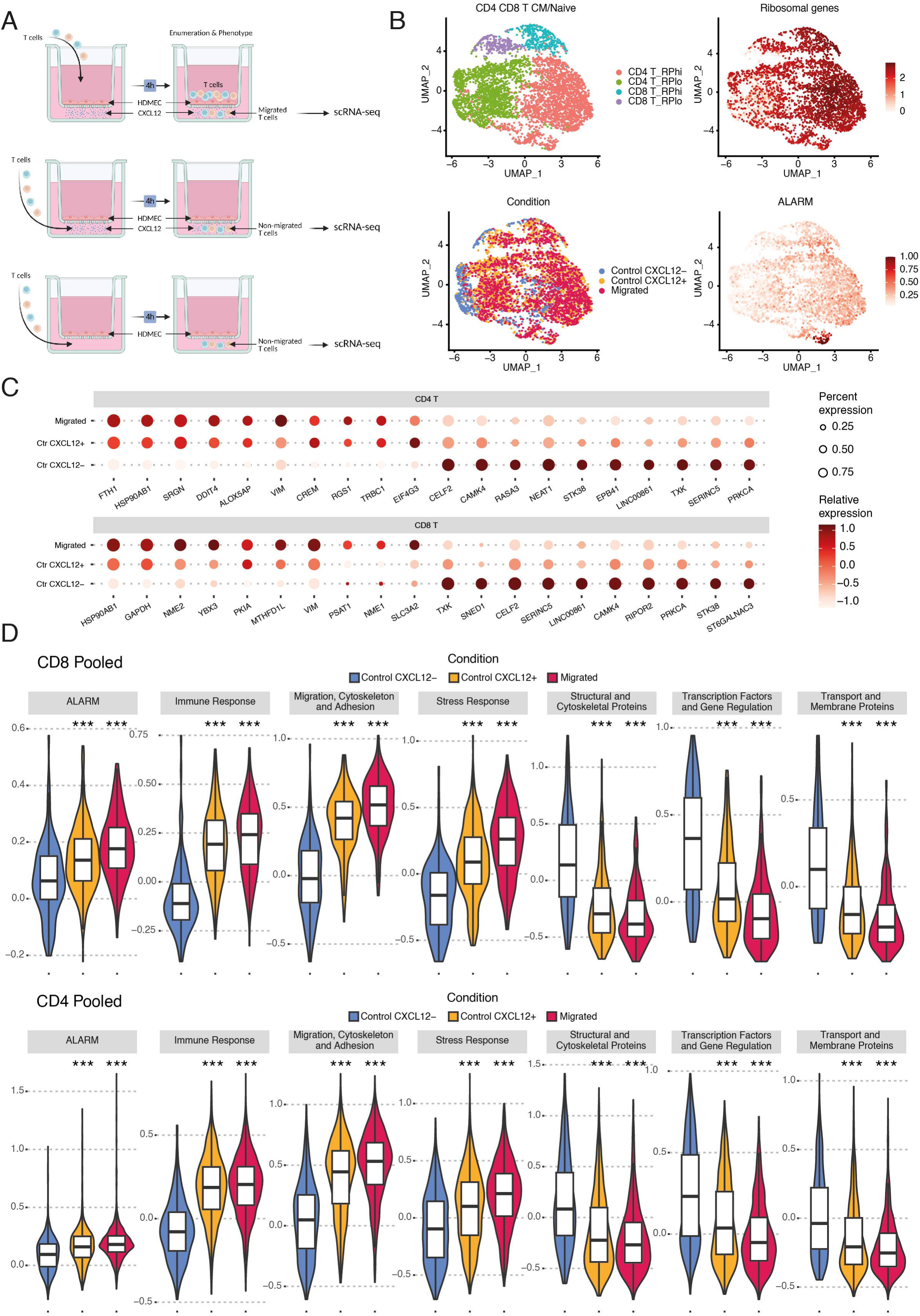
Single Cell Characterization of T Cell Behavior in an In Vitro Transwell Assay. A) Schematic representation of the transwell assay used to investigate CXCL12-mediated signaling and its effect on T cell migration. Three distinct conditions were assessed: (1) CXCL12-: T cells were placed below the transwell membrane without exposure to CXCL12, (2) CXCL12+: T cells were placed below the transwell membrane in direct contact with CXCL12, and (3) Migrated: T cells were placed above the membrane and collected from below after migrating in the presence of CXCL12. B) UMAP visualizations displaying the clustering of naïve CD4 and CD8 T cells based on gene expression. Clusters were identified based on differences in ribosomal gene expression and CD4 versus CD8 annotation (top left & right). Annotation according to condition (bottom left) shows a notable separation between CXCL12-T cells and those exposed to CXCL12 or that had transmigrated. The ALARM module shows lower expression in the CXCL12-condition compared to the CXCL12+ and Migrated groups (bottom right). C) Dot plot showing the expression levels of specific genes across the different conditions (Migrated, CXCL12+, CXCL12-). D) Violin plots depicting the expression distribution of gene groups in pooled CD4 and CD8 T cells across the three conditions. The plots demonstrate changes in gene expression associated with immune response, migration, cytoskeleton organization, and stress response, as well as the downregulation of genes related to structural organization and gene regulation.

To more accurately quantify gene expression changes and minimize the impact of subtype differences, we focused on the most abundant subsets, namely naïve CD4 and CD8 T cells. These naïve T cells clustered into four distinct groups, primarily due to differences in ribosomal gene expression and the annotation of CD4 versus CD8 cells (Figure 6B). Additionally, there was a clear separation between CXCL12-T cells and those exposed to CXCL12 or that had transmigrated, indicating significant underlying gene expression differences. This separation was further evidenced by distinct changes in the expression of ALARM module genes, with the CXCL12-condition showing markedly lower expression patterns compared to the CXCL12+ and transmigrated groups (Figure 6B). Next, we investigated specific gene expression alterations which would increase or decrease from CXCL12-, CXCL12+ to the transmigrated condition (see methods).

In addition to ALARM module genes (Supplemental Figure S4D), several others exhibited notable changes in expression across the different conditions (Figure 6C, Supplemental file 1). HSP90AB1, a member of the HSP90 family of chaperone proteins, which is crucial for stabilizing proteins involved in cell survival and esponses^35^ was significantly upregulated in both CD4 and CD8 T cells. This suggests that HSP90AB1 may play an important role in enhancing the functional stability of proteins required for T cell migration and adaptation during CXCL12 stimulation. Interestingly, two genes with roles in cell migration, VIM^36^ and STK38^37^ (serine/threonine kinase 38), also showed differential expressions in both cell types. VIM, a key regulator of cytoskeletal organization that promotes cell motility^36^, was upregulated in migrated cells, aligning with its role in facilitating the cytoskeletal rearrangements necessary for migration. On the other hand, STK38 was upregulated predominantly in the CXCL12-condition. To better understand the roles of all differentially expressed genes, we grouped them by function (Supplemental file S1, Figure 6D). We found gene sets involved in immune response, migration, cytoskeleton, adhesion, stress response and metabolism were gradually upregulated in the CXCL12+ and migrated conditions in both CD4 and CD8 T cells. Conversely, gene sets associated with structural organization, gene regulation, and membrane transport were downregulated. These findings suggest that CXCL12 signaling, and migration induce profound metabolic and functional changes in T cells, preparing them for new roles that require increased energy and biosynthetic demands. Given the prominent upregulation of metabolic pathways, we further investigated the metabolic reprogramming that accompanies T cell migration and activation in response to CXCL12. To achieve this, we performed a comprehensive metabolic pathway analysis on T cells using Compass, an algorithm designed to characterize the metabolic state of cells by integrating single-cell RNA-Seq data with flux balance analysis^38^. This in silico approach allows us to infer the metabolic status of individual cells based solely on transcriptomic data, providing insights at single-cell resolution. The analysis revealed significant upregulation in several metabolic pathways, notably glycolysis/gluconeogenesis, phosphatidylinositol signaling, and amino acid metabolism in response to CXCL12 and migration (Figure 7A). To visualize the overall metabolic differences between the conditions, we performed PCA on the Compass score matrix, which quantifies the metabolic state in each cell (see methods)^38^. The PCA results showed clear clustering of samples according to their condition, with non-migrated T cells forming a distinct cluster separate from CXCL12+ and migrated groups (Figure 7B). This separation underscores the significant impact of CXCL12-induced migration on the metabolic state of T cells. Because the metabolic shift towards glycolysis is an essential hallmark of T cell activation^39^ we focused on the glycolysis and gluconeogenesis pathways to investigate the gene expression changes involved in these processes (Figure 7C). Key glycolytic enzymes, such as glucose-6-phosphate isomerase and pyruvate dehydrogenase, were significantly upregulated in migrated T cells compared to controls. To validate the transcriptomic findings experimentally, we performed flow cytometry analyses to assess glucose uptake and the expression of glycolytic enzymes. We measured protein levels of glucose transporter 1 (GLUT1) and lactate dehydrogenase A (LDHA), a key glycolytic enzyme that converts pyruvate to lactate for rapid ATP production under anaerobic conditions. Although the increase in GLUT1-positive cells post-transmigration was not statistically significant (p = 0.68) (Figure 7D), LDHA expression showed a significant increase in migrated T cells compared to controls (p = 0.019) (Figure 7D). Additionally, uptake of the fluorescent glucose analog 2-NBDG was significantly elevated in migrated cells (p = 0.031) (Figure 7D), indicating enhanced glucose metabolism. These results confirm an increased glycolytic activity observed in migrated T cells, consistent with the metabolic reprogramming identified in our pathway analysis. In summary, our results demonstrate that T cells migrating in response to CXCL12 undergo functional reorganization, enabling their transition from circulating cells to active immune responders at sites of inflammation or tissue injury.

**Figure 7.**
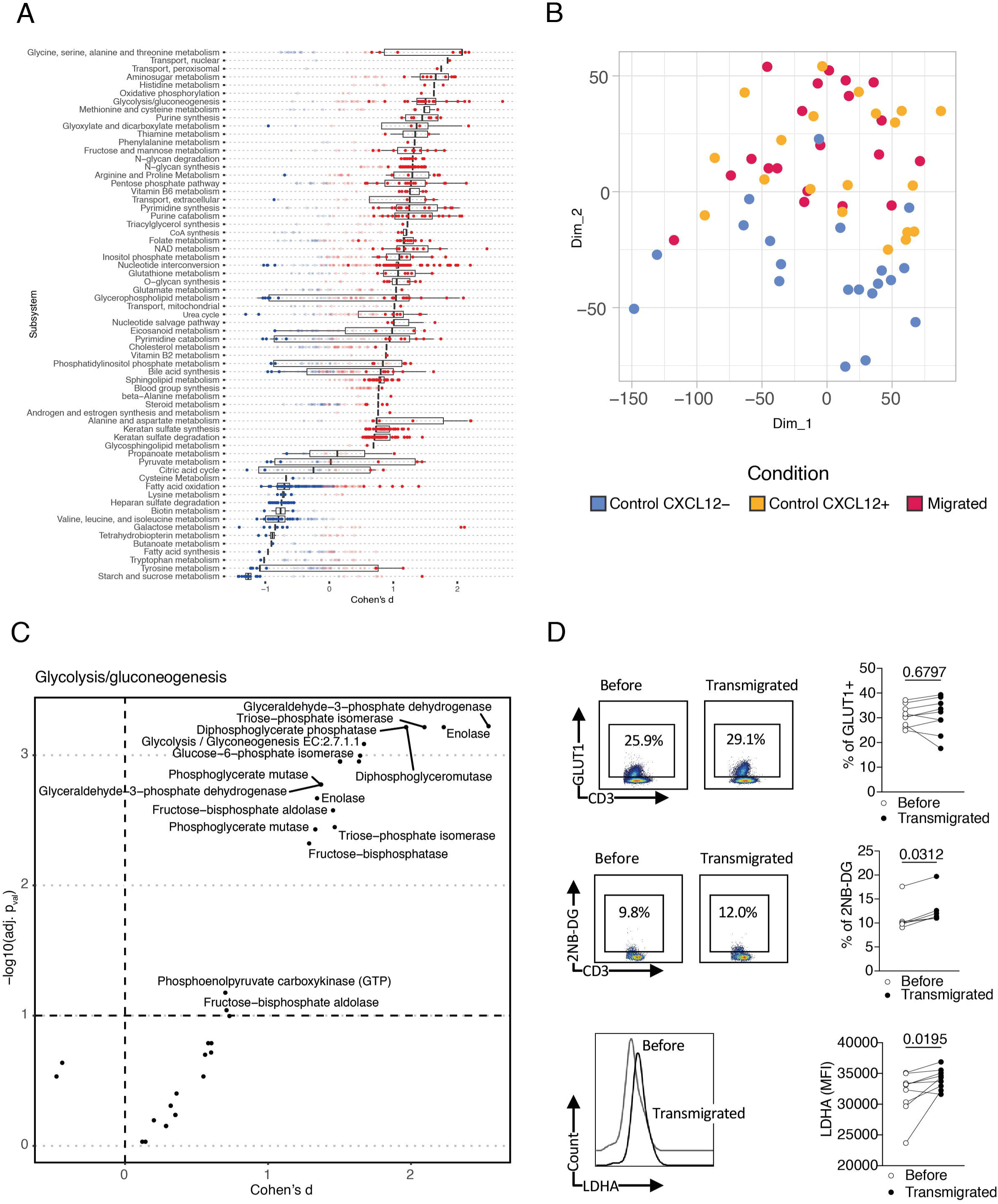
ALARM Module associates with a shift of immune cells metabolic fonctions. A) Metabolome pathway analysis of CD4+ T cells, using 20 pseudobulk sample for each experimental condition (Migrated, CXCL12-positive, and CXCL12-negative). The differential activity of metabolic reactions was evaluated by comparing the mean of Migrated samples and CXCL12-samples B) PCA visualization of the pseudobulk samples by condition based on the metabolome pathway analysis scores C) Volcano plot illustrating glycolytic enzyme changes in migrated T cells compared to controls. The x-axis shows Cohen’s d effect sizes, and the y-axis indicates the statistical significance (−log10 p-values). Key enzymes in the glycolysis/gluconeogenesis pathway are labeled. D) Flow cytometry analysis and quantification of GLUT1 and 2-NBDG uptake in T cells before and after transmigration. Left panels show representative flow cytometry plots, and right panels show paired comparisons for each parameter. LDHA expression levels (measured by MFI) are also shown before and after transmigration. Statistical significance is indicated, with p-values provided for each comparison.

### ALARM is expressed in healthy individuals and variation is associated with infectious disease

Up to this point, we have examined the role of ALARM cells primarily in the context of kidney rejection and stable kidney transplantation patients. However, the findings from the Transwell assay, using healthy donor cells in an in vitro model, suggest that ALARM cells may play a broader role beyond kidney-specific contexts. Therefore, we investigated whether the ALARM module is expressed across a healthy population to assess its broader function. For this, we explored a publicly available scRNA-seq data set of ∼25,000 PBMC from 45 healthy volunteers (HV) for ALARM expression ^40^. The data was generated from the *LifeLines DEEP* cohort in Netherland, ranges in age from 20 to 79 and contained 46,6% female individuals and was described to be healthy time of collection as estimated by two general practitioner visits ^41^. The ALARM gene expression was prominent in all cell types as shown by its module score suggesting that ALARM is generally expressed in HV (Supplemental Figure S5A). We note that age and sex did not result a significantly different expression of the ALARM (Supplemental Figure S5B). Since this module was generally expressed in HV, we next asked whether it may be involved in other disease conditions than transplantation rejection, but which implicate the recruitment of circulating immune cells to specific tissues. To test this hypothesis, we exploited a publicly available scRNA-seq data on PBMCs in which healthy individuals were intravenously injected with the endotoxin lipopolysaccharide (LPS), a component of the cell wall of Gram-negative bacteria ^5^. LPS in the bloodstream causes an immediate systemic release of a variety of inflammatory mediators, a fever and a rapid but transient leukopenia ^42,43^. This experiment is thought to mimic an acute systemic inflammatory response (SIRS) ^5^, and thus provides an ideal proxy of how ALARM expressing cells are responding to LPS-induced SIRS. We used the preprocessed available scRNA-seq data which contained the 0-time point (10 HV), 90 minutes (6 HV) and 10h after the LPS injection (6 HV) and first evaluated how the cellular proportions changed over time. We measured the ALARM score in the three conditions across the CD4, CD8, NK and B cells (Figure 8A). There was rapid and significant reduction of ALARM high expressing cells as soon as 90 minutes and which further decreased until 10h after the LPS injection (Figure 8A). This decrease was continuous within three individuals for which both timepoints were available, indicating that ALARM cells are reduced in a time dependent manner (Figure 8B). This drastic change of ALARM expression in such a short time frame suggests that cells which highly express this module rapidly egress from circulation, directly contributing to the transient leukopenia observed upon LPS injection. This is also consistent with the transient leukopenia associated with LPS i.v. injection. Furthermore, this response to LPS which is thought to be initiated via TLR4 receptor signaling expressed on circulating monocytes ^44,45^ signifies that ALARM is not solely implicated in transplantation rejection or kidney immune cell invasion but appears to also be involved in the inflammatory response to endotoxin.

**Figure 8.**
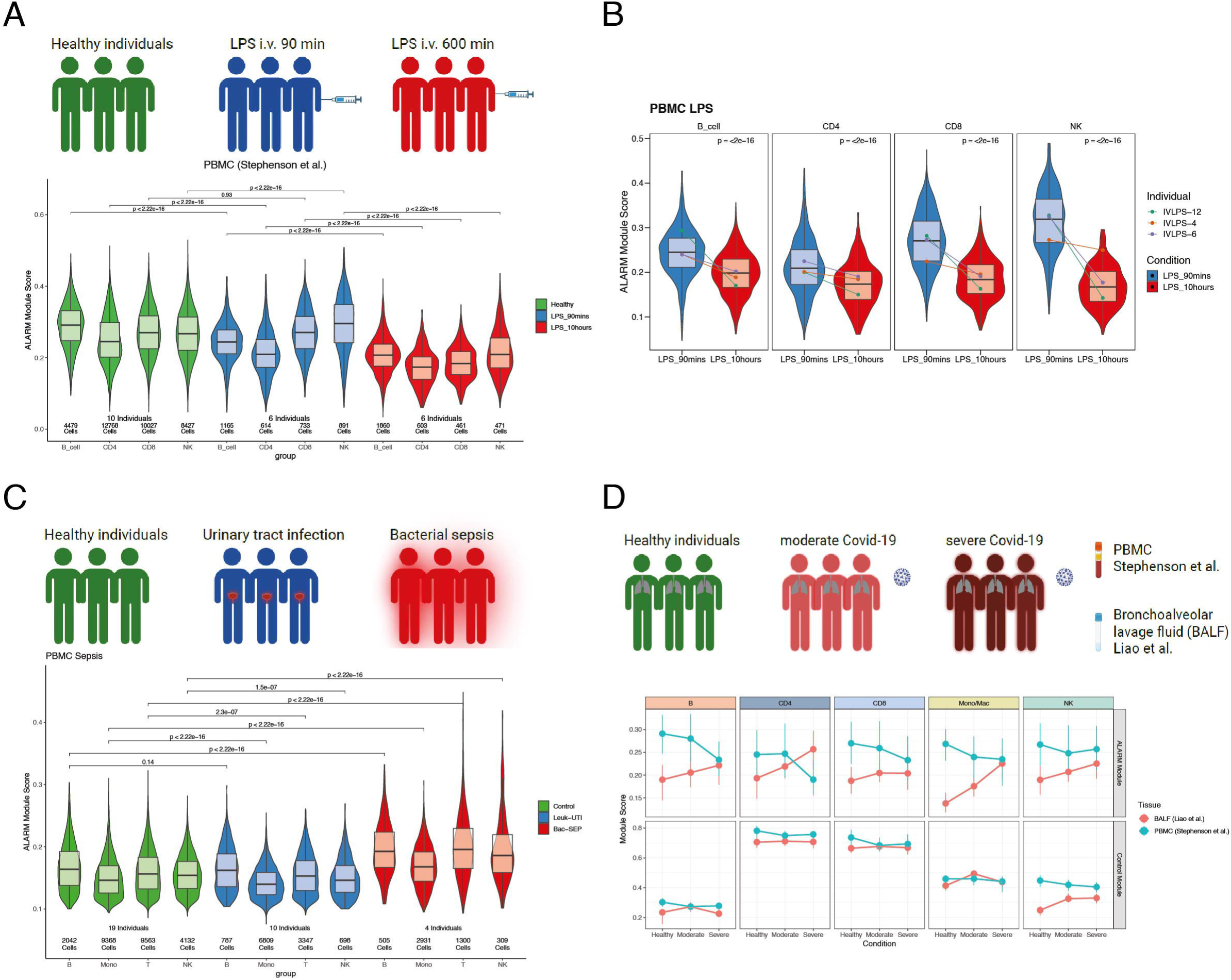
ALARM gene expression is altered in distinct immune conditions. A) Outline of study on Lipopolysaccharide (LPS) intravenous injection (iv) scRNAseq experiment performed on healthy volunteers obtained from Stephenson et al. The annotated expression data was used to compute the ALARM module score across time points after LPS injection in healthy volunteers. Violin plots show the module score across cell types and condition. Total number of cells are mentioned below the plot. P-values are shown above the violin plots and were calculated using Wilcoxon rank sum test comparing each cell type between healthy and LPS conditions. B) Violin plots of ALARM module score for three individuals which had matching timepoints across all cell types in the condition LPS 90 min and LPS 10h. Lines connect the median ALARM module score across time points for each individual separately. P-values are shown above the violin plots and were calculated using Wilcoxon rank sum test comparing the two distinct time points. C) Outline of patients with sepsis and urinary tract infection obtained from Reyes et al,. Annotated data was used to compute ALARM module score across the different cell-types in urinary tract infection (UTI) patients and bacteraemia sepsis patients. Number of cells and individuals used are shown below the violin plots. P-values are shown above the violin plots and were calculated using Wilcoxon rank sum test comparing each cell type between healthy and LPS conditions. D) Outline of PBMC scRNA-seq generated on healthy individuals, moderate covid-19 and severe covid-19 patients (Stephenson et al.,) and Bronchoalveolar lavage fluid (BALF) from a distinct cohort (Liao et al.). Upper panel shows median ALARM module score for each cell-type for PBMC (blue) and BALF (red). Lower panel shows for each cell-type the module score for each corresponding cell-type specific module.

Next, we investigated how ALARM may regulate when the site of inflammation is localized to a single organ as in urinary tract infection. For this we explored a publicly available PBMC scRNA-seq dataset which contained patients with leukocyte infiltrating urinary tract infection (UTI). We chose this condition as the data was generated on patients which presented a localized infection with infiltrating leukocytes. The study also provided results of HV and sepsis patients, notably patients with bacteremia, i.e. bacterial presence in the blood. ^1^ (Figure 8C). The bacteremia patients were thus also used for comparison since this condition reflects a generalized or systemic infection which is distinct from a localized infection such as UTI (Figure 8C). Interestingly, we found that ALARM expression was reduced in circulating cells in UTI patients (except B cells), consistent with the recruitment of leukocytes to the tissue. In contrast, in patients with bacteremia, the ALARM cells accumulated in the circulation, indicating that under this condition ALARM expressing cells may not be recruited to a specific tissue. This contrasted with the response to LPS iv which is also thought to engender a systemic response (see discussion).

To further explore the role of ALARM in response to localized infection, we analyzed two scRNA-seq datasets generated from Covid-19 patients ^5,46^. The aim was to explore the dynamics of ALARM cells in the circulation in comparison to the lung. For this, we compared ALARM expression in PBMCs with ALARM expression in broncho-alveolar lavage fluid (BALF). The two separate original studies stratified the patients by healthy, moderate, and severe Covid-19 disease and we used this stratification to compare the ALARM module expression in the blood (PBMC) and in the lung (BALF) (upper panel figure 8D). The ALARM cells diminished according to disease severity (T, B, Mono, but not NK) in the blood stream. This reduction was concomitant with an increase of ALARM high cells in the lung suggesting that ALARM cells are migrating to the site of infection. Interestingly, these changes were cell-type specific, notably while CD4+ T cells increased it was not the case for CD8+T cells. We note that the cell-type specific modules did not change between disease state and between blood and lung (lower panel, figure 8D) indicating that the similar cell types were analyzed and that the cell-type specific modules were not related to disease state.

Collectively these results show that ALARM displays a normal distribution of expression across healthy individuals and changes in response to distinct disease states (bacterial, viral, kidney rejection). It is noteworthy that expression alterations of this module are apparent in distinct cell types depending on the disease conditions.

### Recruitment of CXCR4+ T Cells During Lung Infection In Vivo

To independently validate the rapid recruitment of immune cells during lung infection, as exemplified in the COVID-19 results above, we used a well-established pneumonia mouse model^47,48^ to study T cell recruitment. The infection was induced with *E. coli* and resolved after 7 days, with the peak of infection occurring between day 1 and day 3. To specifically evaluate the recruitment of T cells from the blood, we employed CD45-PE mediated immune staining of blood immune cells before and during infection^48^ (Figure 9A). This method allowed us to precisely quantify the recruitment of cells from the blood during the infection. We observed that T cells were recruited from the blood as early as day 1, with a significant peak at day 3, indicating a rapid response to the lung infection (Figure 9B). Upon analyzing the membrane expression of CXCR4 on these cells, we found that 15% to 20% were CXCR4+ T cells, a key component of the ALARM module, suggesting that a diverse set of T cells is recruited to the lung (Figure 9B, Pie Charts). Next, we investigated the membrane display of CD69 and found that most CXCR4+ CD69+ T cells were predominantly CD45+ cells (60-80% at day 1 and day 3), indicating their blood-derived origin (Figure 9C). This phenomenon was particularly evident at day 1 and day 3, corresponding with the peak of infection. Additionally, stratification of the CXCR4+ T cells by CD4+ and CD8+ subsets showed that CD4+ T cell recruitment was much more abundant than CD8+ T cells, mirroring observations from COVID-19 lung infection studies (Figure 9D).

**Figure 9.**
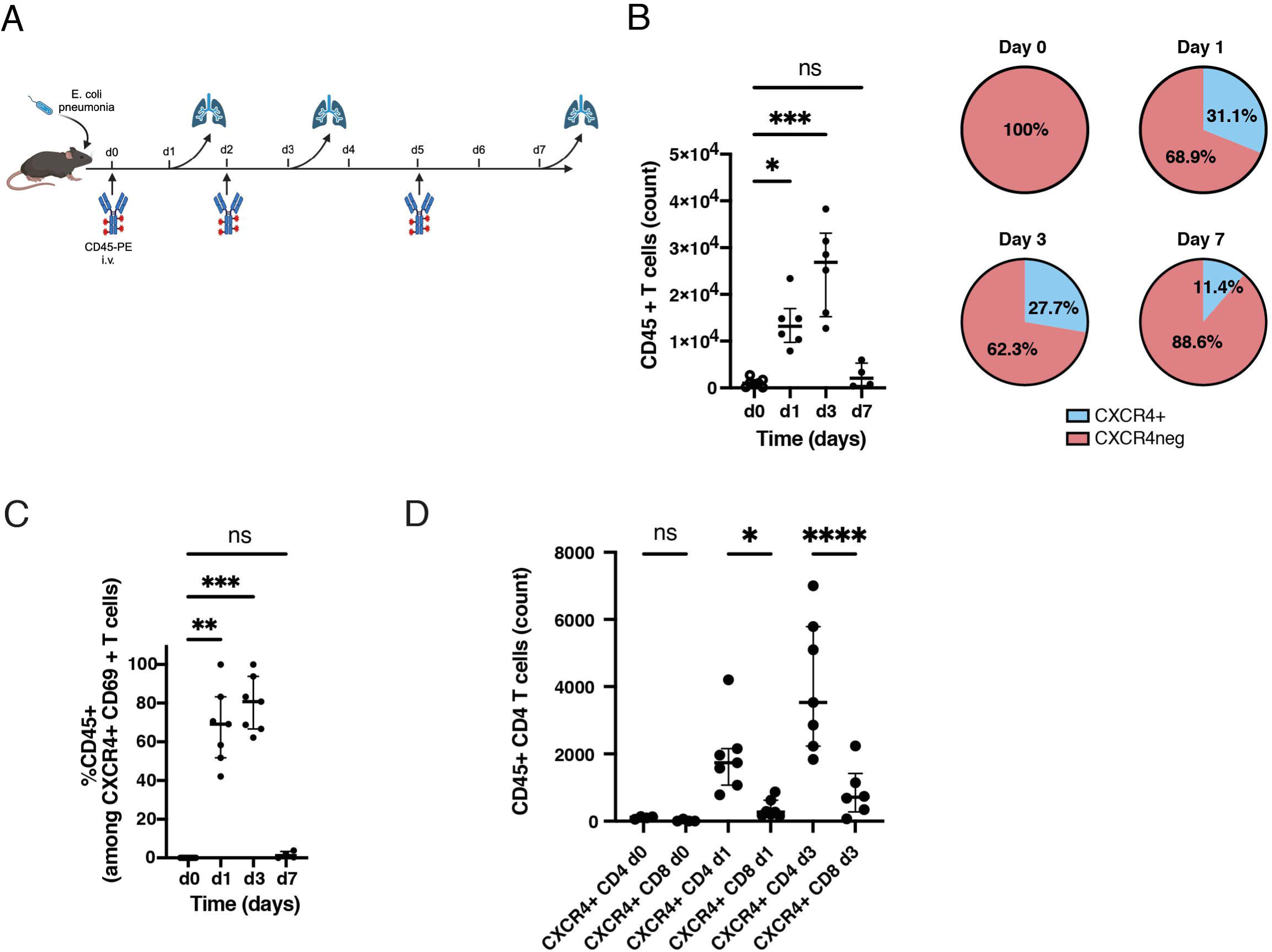
Recruitment and Characterization of CXCR4+ T Cells During Lung Infection in Vivo. A) Schematic representation of the experimental setup used to study T cell recruitment during *E. coli* pneumonia in a mouse model. The infection was induced on day 0, with peak infection occurring between days 1 and 3, and resolution by day 7. CD45-PE was intravenously injected at different time points (days 0, 1, 3, and 5) to stain and track blood-derived immune cells migrating into the lungs during infection. B) Quantification of CD45+ T cells in the lungs at various time points post-infection. The left plot shows the number of CD45+ T cells at each time point. The accompanying pie charts depict the proportion of CXCR4+ and CXCR4-T cells over the course of the infection. C) Percentage of CD45+ cells within the CXCR4+ CD69+ T cell population over time. D) Comparative analysis of CD45+ CXCR4+ CD4+ and CD8+ T cell counts at different time points. Independent experiments with 3-4 mice per condition were carried out, and statistical significance was determined using one-way ANOVA, followed by post-hoc tests where appropriate. Significance levels are indicated as follows: *< 0.05, ** < 0.01, and *** < 0.001.

These findings demonstrate that CXCR4+ T cells, particularly CD4+ T cells, are rapidly recruited from the blood to the lung in response to infection, with peak recruitment occurring between day 1 and day 3. The presence of CD69+ cells among the recruited CXCR4+ T cells suggests that these cells are not only migrating but are also likely being activated or display a residency program^34^. The observation of the concomitant *protein* expression and membrane display of CXCR4 and CD69 is characteristic of the ALARM module, which is defined by the *gene* expression of both CXCR4 and CD69. The results underscore the crucial role of the ALARM module in orchestrating an effective immune response during the peak of lung infection, highlighting its broader relevance beyond kidney-specific contexts.

### Disease classification of ALARM in immune mediated diseases

The above results suggest that ALARM expression changes are associated to immune diseases, likely via the recruitment of circulating immune cells to the site of inflammation. It may thus be possible that the ALARM genes could be used as predictors for immune disease classification. The rationale is that if ALARM genes are relevant for a precise disease state, they would be strong predictors to classify healthy from disease ^49^. To investigate this hypothesis, we analyzed 10 immune-mediated diseases with distinct tissue tropisms (Supplemental Figure S8A). Out of these 10 diseases, systemic lupus erythematosus (SLE), Sjogren syndrome and Anti-neutrophil cytoplasmic antibody-associated vasculitis (ANCA) are known to favor the kidney among other organs. In contrast, mixed connective tissue disease and systemic sclerosis rather favor the connective tissues and rheumatoid arthritis targets the joints. Thus, we aimed to test whether ALARM genes are good classifiers of disease vs healthy condition and whether the classification of disease state varies depending on the tissue tropism. This was possible thanks to a large bulk RNA-seq dataset comprising 28 circulating immune cell-types which included 337 patients across 10 immune mediated diseases and 79 healthy controls ^50^ (Supplemental Figure S8A). We evaluated total gene expression using UMAP of this data set (Figure 10A) and found that ALARM module expression was present in all major cell types (Figure 10B). To test the discriminative ability of the ALARM module, we then devised a classification pipeline comparing disease state (all 10 immune diseases) vs healthy using logistic regression (Figure 10C). As in the data set there were transcriptomics data from 28 cell types available, we focused on B cells, T cells, NK and Monocytes by regrouping their respective sub cell types together (see Figure 10A).

**Figure 10.**
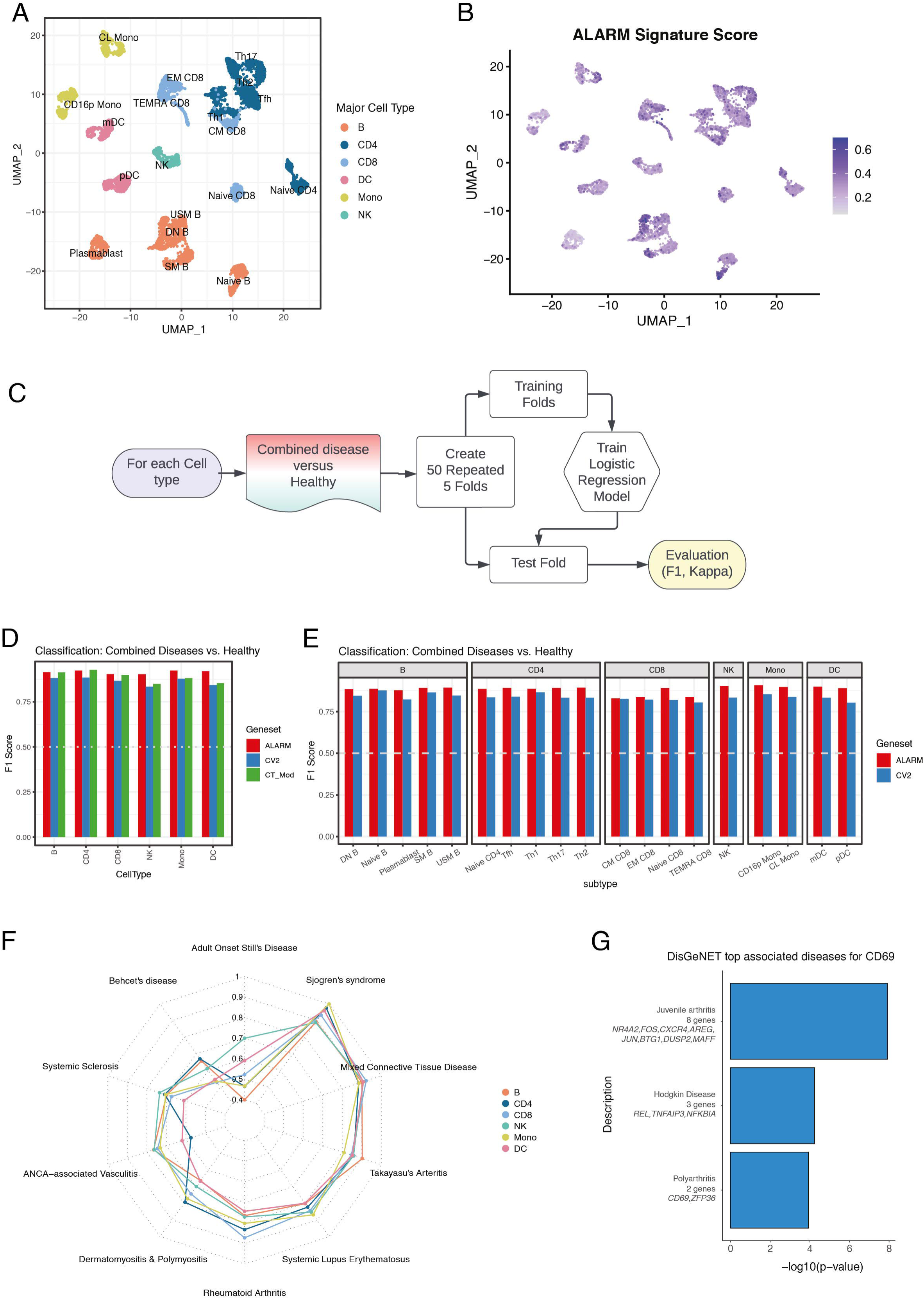
ALARM genes are implicated in and predictive of immune mediated diseases. A) UMAP of bulk transcriptomic study (Ota et al., 2020). Colours represent major cell types and each single point represents a bulk transcriptomic dataset. Cell sub types are annotated in the plot. B) The same UMAP showing expression of ALARM module expression across samples. C) Outline of classification approach used to test disease prediction D) Barchart showing the F-1 score of disease classification for ALARM, cell-type specific modules and coefficient of variation (CV^2^) selected genes in each major cell type separately. E) Barchart of F1-scores computed for disease classification in each sub cell type separately for ALARM and CV^2^ genes F) Radar chart showing the F1-score in each cell type for classification between each disease and healthy separately using ALARM genes in each major cell type. G) Barchart showing the top three enriched diseases in the ALARM genes using the DisGeNET curated database.

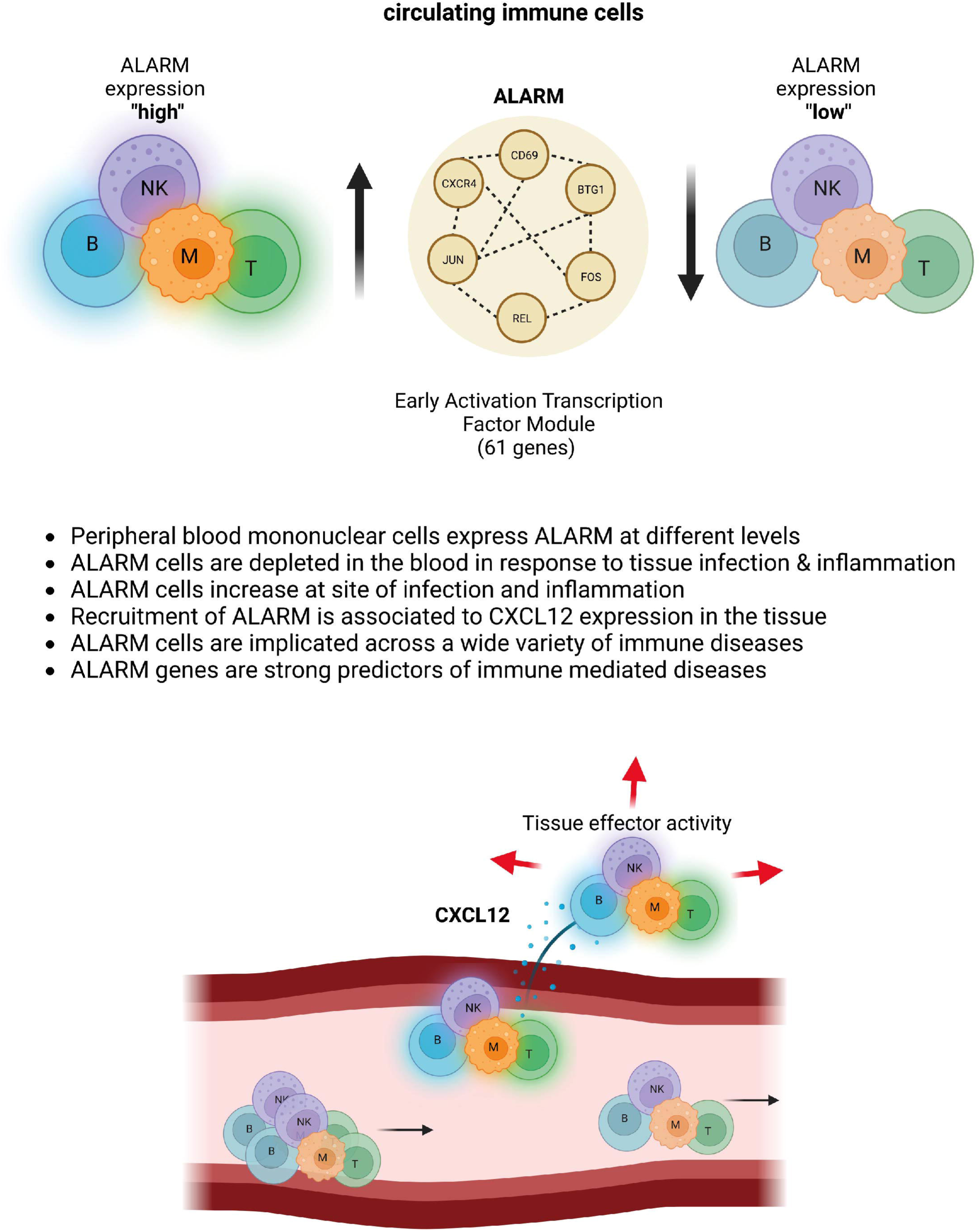

We then generated an ALARM gene classification model compared it to two other models. The first one using the most variable genes (coefficient of variation (CV^2^), see methods) and the second model was based on genes in cell type identity modules (see figure 2). The rationale for using CV^2^ gene selection was to use an independent gene selection process which is more predictive than random gene selection. We compared the prediction performance of the three models using the F-1 score as it gives equal weight to precision and recall (Figure 10D, F1 score). Interestingly, the ALARM genes were the best predictors for CD8, Monocytes and NK cells and were similar in performance to the cell type specific modules of CD4 and B cells. To account for possible imbalances in the numbers of disease and control samples we also computed the Cohen’s Kappa score (Supplemental Figure S8B). The results were consistent with the F-scores. Of note, the CV^2^ gene selection approach was less predictive in all cell types (Figure 8D and Supplemental S8B).

To evaluate whether the ALARM module was prominent for any specific cell sub types within the major cell types (e.g., CD4 T helper cells vs CD4 naïve cells) we estimated disease classification performance of the ALARM module separately for each sub type. Predictability as estimated by F1 score remained robust when each subtype was analyzed separately in comparison to the CV^2^ method (Figure 10E). We note that in certain subtypes there were too few samples to compute an accurate Kappa score (Supplemental Figure S8C). Nevertheless, this result indicates that ALARM was found to be relevant in all the subtypes analyzed.

Next, we evaluated whether each of the 10 immune mediated diseases could be individually classified from healthy (Figure 10F). The best classification ability of ALARM was found for Mixed connective tissue disease, rheumatoid arthritis, Sjogren’s syndrome, SLE and Takayasu’s Arteritis. This indicates that ALARM genes are not specifically predictive for tissue tropism but appear to be relevant independently of the targeted tissue. In most cases ALARM outperformed the CV^2^ feature selection suggesting that ALARM genes are likely to be implicated in their disease etiology. In summary, this comprehensive classification analysis indicates that ALARM genes are strong predictors of disease state across the majority of circulating immune cells and within the 10 immune related diseases.

### ALARM is enriched for genetic disease associations

Since the ALARM genes are strong predictors of immune mediated disease and its general role within multiple cell-types and across multiple infections, immune related and autoimmune diseases, it is likely that ALARM is enriched for genes known to be associated to diseases. To test this, we exploited the DisGeNET database^51^ which provides a comprehensive compilation of published and curated human gene disease associations (GDAs) from repositories including Mendelian, complex and environmental diseases and enables enrichment analysis of such GDAs. Notably, we found that 24 out of the 61 ALARM genes (39%) were associated with a disease. To estimate the probability of this occurring by chance, we compared this to random sampling of 61 genes and their quantification of GDAs (Supplemental Figure S8D permutation). The probability of reaching 40 % of genes or more was below > 0.001, indicating that ALARM genes are highly enriched for GDAs.

Next, we tested whether the ALARM genes were enriched for diseases associations (Figure 10G). The top three disease categories enriched for GDAs were Juvenile arthritis, Hodgkin disease and polyarthritis comprising by themselves distinct 13 genes. This enrichment analysis indicates that ALARM is also genetically connected to disease state. In summary, ALARM is a strong classifier and genetically linked to immune disease and thus is likely to play a general role in multiple immune mediated diseases.

## Discussion

In this study, we gathered a cohort of matching ABMR, TCMR and stable patients and generated a comprehensive scRNA-seq atlas of circulating immune cells across time and conditions. We then identified multiple gene co-expression modules. Five out of nine modules were related to a single cell-type while three were expressed in closely related cells (CD4+, CD8+ and NK cells) and only the ALARM module was prominent in multiple cell-types. The observation that single cell transcriptomes mostly reveal cell-type specific modules was also described by Kotliar et al., ^9^ in which they distinguish between identity (i.e., cell-type specific) gene expression programs (GEP) and activity GEPs. It is possible that cell-type specific co-expressed genes are better detected as they show a greater coherence within a well-defined group of cells. This is also notable in the presented data as the gene expression scores of cell-type specific modules show less variance than the ALARM module. Nevertheless, cell-type specific gene expression may not necessarily imply that it remains constant across conditions. For example, we noted that some cell type specific modules were associated to disease state (Figure 2E), notably the NK cell and monocyte specific modules were increased during rejection. It is also possible that because of the relatively low number of genes per cell detected when compared to bulk transcriptomics, cell identity programs are preferentially detected, and more subtle condition specific modules are not robustly detected. Indeed, while cNMF revealed additional modules in the separate batches, only the ALARM module was consistently identified across the three batches. The ALARM genes were found to be highly enriched for transcription factors and gene ontology pathways associated with the gene expression machinery including transcription, mRNA processing and ubiquitination. Prominent transcription factors included the AP-1 complex and the NFKB subunit REL which are both associated with stress responses and inflammation. The membership of CD69 in the ALARM module also suggests a role of stress response. CD69 is a classical early activation marker of lymphocytes, as demonstrated by its rapid display on the surface of T cells after TCR stimulation^34^. In addition, CD69 is also known to be a tissue retention marker as it is expressed on resident memory T cells in distinct tissues. In blood, this gene has been associated with chronic inflammation in various diseases including rheumatoid arthritis ^52^ and systemic lupus erythematosus ^53^. Concomitant with this, CD69 protein expression is increased on infiltrated immune cells at the site of inflammation in immune mediated diseases including systemic sclerosis, rheumatoid arthritis, systemic lupus erythematosus^34^. The membership of CD69 in this module thus indicates that this module role could be to prepare circulating cells T cells for TCR stimulation and for tissue retention once moved into a tissue, i.e., to become T resident effector cells. This notion is also consistent with the increased expression of the ALARM module in the kidney biopsy transcriptomics data (Figure 4).

The cytokine receptor CXCR4 was also identified in this module. CXCR4 is predominantly expressed by lymphocytes as well as monocytes and through which the CXCL12 ligand promotes chemotaxis to tissues via a concentration gradient ^54^. CXCL12 is expressed in multiple tissues including the kidney and is altered during pathophysiological responses including immune diseases. Indeed, an alteration of CXCL12 expression was observed in the kidney transplantation biopsies and this increase was associated to elevated ALARM gene expression in the tissue (Figure 4E). It is possible that the CXCR4-CXCL12 axis contributes to the recruitment of ALARM expressing cells in the case of kidney transplantation rejection and other immune diseases. This is also consistent with the observation that cells expressing the ALARM module decrease in the circulation during kidney graft rejection. This observation was confirmed by both transcriptomics and histological studies of pigs as well as transcriptomics in human kidney biopsies, thus, via CXCR4-CXCL12 leading ALARM cells to infiltrate the tissues during rejection. This mechanism was further supported by an in vitro trans well assay, where CXCL12 was shown to induce T cell migration. More importantly, it was found that both CXCL12 presence and migration significantly increased CD69 protein expression at the cell membrane. Specifically, the combination of HMEC contact and CXCL12 presence was necessary for the highest expression of CD69, similar to that observed during transmigration. Single cell transcriptomic analysis of migrated and non-migrated cells further revealed that certain gene groups, particularly those involved in cytoskeleton organization, migration, and immune response, were upregulated, while others were downregulated, indicating a shift in cellular state to adapt to new functions after migration. Furthermore,we found that migration of T cells in response to CXCL12 is accompanied by significant metabolic reprogramming. We observed upregulation of glycolytic pathways, increased expression of LDHA at protein levels, and enhanced glucose uptake in migrated T cells. Metabolic reprogramming towards increased glycolysis is a hallmark of activated T cells and is essential for their effector functions during immune responses ^39^. These findings suggest that ALARM module expression not only primes T cells for migration and tissue retention but also prepares them metabolically for the demands of their new functional roles at sites of inflammation.

This CXCR4-CXCL12 axis also highlights the notion that the ALARM module is not necessarily specific to transplantation rejection or the kidney. Indeed, reanalysis of circulating immune cells from publicly available scRNA-seq data showed that the ALARM module was expressed in 45 unrelated healthy individuals ^40^ and showed significant alteration between distinct pathological conditions (Figure 5). First, the ALARM response to LPS iv injection in healthy individuals revealed that it is time-dependent, illustrated by a gradual decrease of ALARM cells within the same individuals over time. Second, there was a significant difference between ALARM cells depending on the location of the pathological condition. ALARM cells were shown to be decreased in response to kidney rejection, in response to leukocyte infiltrating UTI and Covid-19 infection of the lung highlighting the role of ALARM in the recruitment of cells to the site of inflammation and infection. This was further supported by bacteremia sepsis, a state of systemic inflammation in which ALARM cells were increased in the blood. While LPS iv injection and bacteremia induce both a systemic immune response, the former is associated is essentially an endotoxemia response associated to a transient leukopenia^42,43^ while the latter is a complex and heterogeneous condition that involves multiple factors beyond LPS, such as pathogen virulence, host susceptibility, and coexisting medical conditions. It thus makes sense that leukopenia is associated to the decrease of ALARM, while in bacteremia ALARM expression is increased. Third, ALARM cells showed a gradually measurable response to disease severity. This notion was observed by combining two distinct and complementary Covid-19 datasets one of which was collected on BALF, and which had stratified their patients according to disease severity. ALARM cells decreased in response to severity in the blood with a corresponding increase in the lung.

Fourth, we collected several lines of evidence suggesting that ALARM cells are indeed recruited to the site of inflammation and/or infection. During acute rejection induced in the pig model there was a rapid infiltration of leukocytes concomitant with the reduction of ALARM cells in the blood. The analysis of kidney biopsies revealed an increase of ALARM gene expression during kidney transplant rejection. Similarly, the recruitment of ALARM cells to the lung was observed during Covid-19 lung infection. Finally, an in vivo mouse model of *E. coli* pneumonia demonstrated that CXCR4+ and CD69+ T cells are rapidly recruited from the blood to the lung during the peak of infection further supporting the role of the ALARM module in mediating immune cell recruitment to sites of inflammation.

We thus propose a model in which ALARM expression priorities the infiltration capacity of each circulating cell (see figure 11 model). This model has wide ranging consequences in precision medicine as blocking of ALARM cells to the site of inflammation in the case of kidney rejection may prevent further organ damage or attenuate the immune response in the case of Covid-19 lung infection. It may also be useful to predict disease state as we have shown in figure 8. ALARM was a strong classifier of immune disease when compared to healthy individuals. The importance of ALARM was independently demonstrated by its enrichment for genes mutated in notably juvenile and poly-arthritis. Further investigation is however required to test the specificity of disease detection or whether ALARM is merely a response to inflammatory state in the circulation.

There are several limitations of this study, first it is based on gene transcription and thus remains to be explored for protein expression, however this is difficult to achieve at single cell resolution and we are not aware of any gene co-expression modules estimated at the protein level. Nevertheless, the transwell assay and in vivo mouse model experiments indicate a connection between the CXCL12-CXCR4 axis and CD69 and their display during and after migration. While direct evidence of the recruitment of these cells to the tissue has been experimentally confirmed the module expression was shown to be low in the blood and high in the kidney tissue during rejection which may also be caused be lack of the source of these cells, or a slowing in cellular maturation before expressing ALARM genes an avenue that should be explored in subsequent studies. A further limitation is that as of now we do not have a protein surface marker panel that could be associated to cells with high or low ALARM expression. Such markers would enable the purification of ALARM cells enabling further molecular and cellular characterization. Markers would also allow the targeted modulation of the recruitment of ALARM cells to the graft or during Covid19 as well as other immune mediated diseases may thus impact disease severity. Nevertheless, our study remains important in terms of precision medicine, highlighting the discovery of ALARM, which expression enables cells to be preferentially recruited to the inflamed tissue. This notion is likely to open novel strategies of disease monitoring and disease intervention.

## Methods

### Study of kidney transplant patients Kidney transplantation patients

The PBMC samples used in this study (see table 1) were obtained from the DIVAT biocollection (CNIL agreement n°891735, Réseau DIVAT: 10.16.618). Every patient included in the study was enrolled in the DIVAT biocollection following their informed consent. The PBMC from patients were isolated from kidney transplantation biopsies, frozen with DMSO 10% and stored in liquid nitrogen at the Centre de Ressources Biologiques (CRB, CHU Nantes, France).

### Cell preparation

Frozen PBMC samples were rapidly thawed and resuspended in complete Roswell Park Memorial Institute (RPMI) 1640 media (Invitrogen, Carlsbad, CA) with 5% FBS, pre-heated at 37°C.

Following washing steps in PBS+0.04%BSA, cell pellets were resuspended in 200µL FACS buffer (1X PBS supplemented with 2mM EDTA, 2% FBS) in which dead cells were labelled by adding 0.1 μg/mL DAPI (Invitrogen, Carlsbad, CA). Cells were filtered on 70 μm cell strainer and living cells were then sorted using a Fluorescence-activated cell sorting (FACS) Aria II cell sorter (BD Biosciences, Mountain View, CA).We used the same method as previously described for single cell RNAseq^55,56^. One million cells were kept for each sample and resuspended in 100µL of staining buffer (PBS,2%BSA,0.01% Tween) according to the cell hashing protocol^13^ recommendations. Cells were incubated for 10 min with 10µL of human Fc blocking reagent. Each sample was then mixed with 1uL of a specific TotalSeq-A hashtag antibody (BioLegend, San Diego, CA) and incubated on ice for 30 min. Following 3 washing steps with the staining buffer, cells were counted and their viability measured using an ADAM-MC automatic cell counter (NanoEntek, Seoul,South Korea) to ensure a viability above the recommended 70%. All the samples were pooled at an equal cell concentration in a single vial, centrifugated and resuspended in PBS to obtain a concentration of 700 cells/µL, to match the targeted cell recovery of 32,000 cells. Encapsulation of single cells was performed on a 10XChromium (10X Genomics, San Francisco, CA) with the Chromium Single Cell 3′ Library and Next GEM reagent kit v3. The libraries were sequenced twice for each of the three experiments on the NovaSeq 6000 (Illumina, San Diego, CA) with S1 flow cells. The sequenced libraries were aligned to the GRCh38-2020-A reference genome with CellRanger v5.0.0 (10X Genomics, San Francisco, CA). The scRNA-seq was performed in 3 different experiments following the same protocol. Each experiment included longitudinal samples from three patients (one stable patient, one humoral rejection and one cellular rejection) as well as one late sample of a tolerant patient.

### Method demultiplexing and Seurat analysis

The count matrices were analyzed in R 4.0.3 using the Seurat R package (v4.0.2, Satija Lab^14^). Each experiment was first processed separately, with the same workflow. First, following the standard workflow recommendations, cells with less than 200 unique feature counts were removed (potential empty droplets). Cells with a percentage of mitochondrial genes greater than 15% were excluded as it results from mitochondria degradation from dead or dying cells. The hashing antibody sequences were then collected to demultiplex and assign each cell to its sample using the MULTIseqDemux function. Cells with too little labels information were called “Negative” while cells with a high count of two or more different oligo-conjugated antibody sequences were called “Doublets”. Only cells with a unique HTO were kept for downstream analysis. Singlet cells were annotated automatically with the Azimuth workflow within Seurat, by mapping the query cells on an annotated reference of 162,000 PBMC measured with 228 antibodies^15^.

All the runs were then merged in a single Seurat object. Doublets and contaminant cells to exclude were selected by identifying cells co-expressing marker genes from distinct cell types. After normalization of the global object, the 2,000 most variable genes in the data were selected to compute the correction using the reciprocal principal component analysis (RPCA). The final annotated and corrected object gathering the 12 patients was composed of 50,507 cells.

### Gene Module Identification

Consensus Non-Negative Matrix Factorization (cNMF)^9^ was used to decompose the cell vs gene expression matrix into cell vs module and usage vs gene low-rank matrices. Non-Negative Matrix Factorization is a stochastic method and therefore it was run with 200 NMF replicates to find a consensus robust factorization. For each batch, the top 2000 over dispersed genes were selected as input to the cNMF run. Different K values (7 to 14) were explored to determine the optimal number of modules. For each K, the *stability* and *error* metrics were examined. The best K was chosen such that the error was minimum, and the stability was maximum. Each batch was independently processed to mitigate batch effects from the three different runs. For the three batches, the optimal number of modules (K) were 14, 11 and 13.

#### Module filtering

All modules from the three batches were collated and hierarchical clustering was performed to identify matching modules. Jaccard similarity score was used to define the similarity between two modules. Finally, only those clusters were retained that could represent all three batches. In this way, nine consistent modules were identified. A unique geneset was determined for each consistent cluster by intersecting the top-ranking 200 genes from the modules. The threshold was achieved by observing a scree plot of input number of genes vs the number of genes after intersection.

#### Module association

The genes in the module were examined for a module to be associated with a known cell type. The modules containing marker genes were associated with their respective cell type; for example, the module with MS4A1 and CD79A genes was associated with B cells. However, one of the modules (Module 9, later named as the ALARM module) could not be associated with a known cell type as it did not contain cell type specific markers and was well expressed in multiple cell types.

### UCell module score

The enrichment of a particular set of genes in an individual cell was measured with the UCell^58^ module score. The score is calculated using the Mann-Whitney U statistic, which compares the expression levels of the module genes relative to the total gene expression of the cell. The U-statistic outcome is then normalized between 0 to 1 to produce the UCell score.

### Regression analysis

Change in the module score (µ) along time (τ) for each condition was modelled with linear regression method. The ‘lm()’ function from R *stats* package was used to fit a distinct model µ ∼ τ, per celltype within the group of Stable, ABMR and TCMR individuals. A positive slope indicated that the module score increased with time.

### Gene Ontology analysis

The gene ontology (GO) functional enrichment of the ALARM module gene list was performed using the R package WebGestaltR (v0.4.4) for Biological Process (BP) and Molecular function (MF) annotation. P-value are obtained with the hypergeometric test for ORA (Over-Representation Analysis). As background dataset for enrichment the top 2000 variable genes from each run were merged and used.

### Transcription factor enrichment analysis

To test for enrichment of transcription factors in the ALARM module, GSEA molecular signature database was used to count the transcription factors gene family in the module and in the background dataset used for gene ontology analysis. Fisher exact test was used to calculate the P-value and enrichment of transcription factors in the ALARM module.

### Allogeneic kidney rejection model in pig Animal model

The study protocol was approved by the French Ministry of Higher Education, Research and Innovation (APAFiS #30136). The experiment was performed on 60 to 80kg male pigs (*Sus scrofa*). Test card with pre-applied antibodies from Serafol (Berlin, Germany) were used to identify the pigs’ ABO blood groups. The alloreactivity was performed by mixed lymphocyte reaction assay between donor and recipient. The donor pig was selected from a different breed as inbred pigs might escape rejection. Donor and recipients blood groups were ensured to be compatible to avoid hyper-acute rejection, and mixed lymphocyte reaction assays were positive thus proving their alloreactivity.

### Allogeneic transplantation

Unilateral nephrectomies were performed on two recipient pigs under general anesthesia with a premedication by Zolazepam/Tiletamine (Zoletil ® Virbac, Carros, France) 15 mg/kg IM, before intubation and a maintained ventilation with a mixture of 49% oxygen, 49% nitrous oxide and 2% isoflurane. The two kidneys from a third donor pig were harvested in the same operating time. The two recipient pigs received one collected kidney each for an orthotopic transplantation. During surgery, a central venous catheter (CVC) was inserted into the internal jugular vein for hydration and medication. Post-operative analgesia was performed every day with intravenous injections of Nalbuphine (Nubain ®, Mylan, Canonsburg, Pennsylvania) and Paracetamol at a dose of 25 mg/kg. Prophylactic antimicrobial therapy was conducted with Cefazolin 1 g (Cefovet ®, Dopharma, Ancenis, France).

### PBMC collection in Pig model

Blood samples were collected daily through the CVC and frozen in a CoolCell® container (Corning ®, Corning, NY, USA) at −80°C following the PBMC isolation. Kidney transplant biopsies were collected daily using automated biopsy needles of 16 gauges under ultrasound guidance while pigs were sedated by Zolazepam/Tiletamine and locally anesthetized with Lidocaine. Kidney samples were then placed in cryovials with 1mL fetal bovine serum (FBS) and 10% dimethylsulfoxyde (DMSO) for gradual cooling in a CoolCell chamber.

### Single cell preparation for pig model

Blood samples were processed as described previously (see Methods 1.2). After filtering, cells were centrifugated at 300g for 10 minutes at 4°C and resuspended in staining buffer for the HTO antibody labeling (see Methods 1.3). The sequenced libraries were aligned to the Sscrofa 11.1 (February 2017 release) reference genome with CellRanger v5.0.0.

### Pig biopsy immunostaining analysis

Kidney biopsies fixed Carnoy’s solution for 30 minutes followed by a fixation in formaldehyde for 24h for optical microscopy purpose. A second batch of kidney biopsies was prepared for immunostaining purpose: biopsies were placed in cryomold, covered with optimal cutting temperature (OCT) compound and immersed in cold isopentane. Following their solidification, cryomolds were stored in liquid nitrogen. Cryosectioning was performed and the resulting slides of kidney biopsies were stained with periodic acid-Schiff (PAS) and Masson’s trichrome stains (TM).

The cellular infiltration was counted using ImageJ^59^ software on the PAS-stained kidney biopsies. Areas of interest were selected to exclude areas with glomeruli. Pictures were first converted to 8-bit grayscale, and the threshold of detection was set to capture only the stained cells.

### Transmigration Model

HDMECs (10 × 10^4^ cells) activated with TNF-*α* (100 U/ml) for 24 hours were seeded O/N onto 1% gelatin-coated Transwell membrane inserts (24-well, 3-*μ*m-pore polycarbonate membrane; Corning Life Science) in endothelial cell growth medium at 37°C. On the day of the assay, purified CD3 T-cell subsets (4×10^5^) were added to the upper transwell migration chamber, and the chemokine CXCL12 (50 ng/ml) was added to the lower transwell migration chamber. Migration was assessed after 4 hours by quantifying the number and phenotype of migrated cells in the lower chamber using 123count eBeads counting beads and a Cytek AURORA flow cytometer (5 lasers). Migrated CD3 were surface stained with specific antibodies to characterize phenotype CD3, CD8, CD4, CD45RA, CCR7 and activation molecules CD69, CD25, CD127, CD95, CD103 and CD49. The antibodies used for the cytometric analyses are listed below <u>Expression of Cytotoxic Molecules by Human transmigrated CD3 T Cell</u> <u>Subsets</u>

To define the expression of cytotoxic molecules, transmigrated CD3 were restimulated with PMA (50ng/mL), ionomycin (500ng/mL) and BFA (5ug/mL). Transmigrated CD3 were surface stained with specific antibodies for phenotypic characterization of CD3, CD8, CD4, CD45RA, CCR7, and after fixation and permeabilization (BD Cytofix/Cytoperm), intracellular staining was performed using antibodies against granzyme B (GZMB) and perforin-1 (PERF-1), granulysin, and TNFa. The antibodies used for cytometric analyses are listed below.

### Metabolic characterization of Human transmigrated CD3 T Cell Subsets

Transmigrated T cells were stimulated over-night with plate bound anti-CD3 (1ug/mL) and anti-CD28 (2ug/mL) mAb. Cells were washed, surface stained with anti-CD3, CD4 and CD8 mAbs and cultured for 30’ at 37°C 5% CO2 in glucose-free medium containing 50 μM 2-NBDG. Alternatively, cells were surface stained with anti-CD3, CD4, CD8 and GLUT1 mAbs and after fixation and permeabilization (BD Cytofix/Cytoperm), intracellular stained with anti-LDHA mAb. Data were acquired using a 5 lasers Cytek AURORA flow cytometer and analyzed using OMIQ.

Antibodies used:

| Target | Clone | Fluorochrome | Provider |
| --- | --- | --- | --- |
| CD3 | OKT3 | Purified | In house production |
| CD28 | CD28.2 | Purified | In house production |
| CD3 | SK7 | PE Cy7 | BD |
| CD4 | SK3 R7 | CF568 | Cytex |
| CD8 | SK1 | Sparkblue 550 | Biolegend |
| CD69 | FN50 | APC Cy7 | Biolegend |
| GLUT1 | 202915 | FITC | R&D Systems |
| LDHA | APC | ab310866 | Abcam |
| CD3 | UCHT1 | PE-Cy7 | Biolegend |
| CD8 | SK1 | Spark Blue 550 | Biolegend |
| CD4 | SK3 | cL-Fluor YG584 | Cytex |
| CD45RA | HI100 | BUV496, BV786 | BD |
| CCR7 | G043H7 | APC Fire810 | Biolegend |
| CD69 | FN50 | APC-Cy7 | Biolegend |
| CD25 | M-A251 | PE-Fire700 | Biolegend |
| CD127 | HIL-7R-M21 | APC R700 | BD Biosciences |
| CD95 | DX2 | PE-Cy5 | BD Biosciences |
| CD103 | Ber-Act8 | BV605 | BD Biosciences |
| CD49a | SR84 | BUV661 | BD Biosciences |
| GZMb | QA16A02 | PE-Cy5 | Biolegend |
| IFNg | B27 | APC | BD Biosciences |
| GNLY | RB1 | AF488 | BD Biosciences |
| TNFa | Mab11 | BUV737 | ThermoFisher |

### scRNA-seq of Transmigration model

T cells were collected as indicated in figure 5A and then processed for chromium loading. The three conditions were processed at the same time using CITE-seq approach (see above).

#### Primary Analysis

Fastq files were generated from Illumina bcl files using Bcl2fastq version 2.2. Cellranger v7.2 was employed to create a filtered scRNA gene expression matrix from the fastq files, utilizing default parameters and the human genome reference version GRCh38-2020.

Seurat version 5.1 was used for subsequent quality control and preprocessing. HTODemux, with a positive quality threshold of 0.95, was applied to demultiplex cells, identifying singlets and associating each singlet with the corresponding condition. Cells with an RNA count exceeding 30,000 or exhibiting mitochondrial gene expression above 20% were excluded. Cell type annotation was performed using Celltypist annotation tool^60^.

#### Secondary Analysis

For each CD8 and CD4 cell type, genes exhibiting significant gradients across the three conditions were identified using a linear model. The dependent variable in the model represented the condition, with control CXCL12-, control CXCL12+, and migrated groups assigned values of 0, 1, and 2, respectively. To simulate multiple individuals, cells were randomly grouped into 10 groups. These groups served as pseudo-individuals, each containing at least 50 cells, created using the createfolds function from the caret package. The final linear model was formulated as ∼ condition + (1 | individual). Top genes were identified based on significant p-values (< 0.05), and the direction of their gradients across the conditions was noted.

Among the top gene modules, several were found to participate in known pathways. The average gene expression for these modules was calculated using the AddModuleScore function of Seurat.

#### Metabolome Pathway Analysis

The Compass algorithm^38^ was used to characterize the metabolic states of CD4+ T cells across three different conditions: Migrated, CXCL12-positive (CXCL12+), and CXCL12-negative (CXCL12-). The algorithm designed to infer the metabolic state of cells from scRNA-Seq data through flux balance analysis. It addresses the limitations of traditional metabolic assays in assessing metabolic states at the single-cell level, leveraging transcriptome data to predict metabolic activities. For reference to metabolic reactions and pathways, the RECON2 database was utilized for this analysis. The scRNA data were first micropooled, resulting in 20 pseudobulk samples for each condition in order to compare the same number of samples between conditions. The reaction penalties were estimated for various metabolic pathways based on gene expression levels in each pseudobulk sample. Reaction penalties were then converted to negative log scores, with higher values indicating greater predicted activity. Significant active reactions were identified using the Wilcoxon rank-sum test, comparing the Migrated samples to the CXCL12-samples. Reactions with an adjusted p-value of less than 0.1 were considered significant.

### Analysis of the ALARM module in kidney biopsies

Two separate studies were analyzed here. The Reeve *et al.* Affymetrix Microarray data in RAW CEL format was downloaded from Gene Expression Omnibus (GEO) website with accession number <u>GSE98320</u>. The samples were pre-processed using robust multi-chip averaging (RMA) implemented in Bioconductor. The patient condition was obtained from the ‘d96’ metadata column as designated in the corresponding Series Matrix file. The patients strictly defined as either TCMR (n=76), ABMR (n=197), Mixed (n=39) or no major abnormality (STA, n=257) as a stable condition, were retained for downstream analysis. The Callemeyn *et al.* dataset was downloaded from GEO with accession number <u>GSE147089</u>. The CEL files were similarly pre-processed using the RMA method. The labels for each sample were obtained from the Series Matrix file and the phenotypes are defined as biopsies without ABMR (n=168), DSA negative ABMR (n=26) and DSA positive ABMR (n=30). The score used to stratify patients was computed by averaging the z-scores of the ALARM module genes.

### ALARM mean z-score distribution across conditions

The Kolmogorov-Smirnov (KS) test was then used to compare the distribution of the ALARM module score between stable and rejection conditions. This non-parametric statistical test compares the cumulative distribution functions of the mean ALARM z-scores in both groups. The KS test statistic (D) is the maximum vertical distance between the two distributions. The p-value of the test is the probability of obtaining a test statistic as extreme as D or more extreme, assuming that the null hypothesis is true. The null hypothesis is that the two samples are drawn from the same distribution.

### Receptor-Ligand analysis

The ‘iTalk’ R package was used for the receptor-ligand (RL) analysis. The receptors were gene candidates in single-cell kidney transplant stable and rejection patients. The *rawParse()* function with stats=’mean’ was used to identify the candidate genes. For each celltype, the genes were ordered by their average count expression. Only the top 50 percent of these genes were selected for the subsequent RL analysis. The same criteria were adopted for selecting the ligand candidates from the bulk RNAseq biopsy data. The significant interaction pairs were discovered from the iTalk database restricted on the cytokine interactions only.

### Analysis of publicly available scRNA-seq datasets

#### PBMCs of 45 healthy Volunteers (Van Der Wijst MG et al.)

Processed (de-anonymized) single-cell RNA-seq data and its relevant meta data was obtained from the European Genome-phenome Archive (EGA) accession number <u>EGAS00001002560</u>. The available data was merged to build a single Seurat object for downstream analysis. Azimuth reference for PBMCs was used to annotate the cells.

#### LPS and Covid PBMC dataset (Stephenson et al.)

The processed data was downloaded from Array Express under accession number <u>E-MTAB-10026</u>. Only individuals from the same batch containing the LPS-treated volunteers were selected to mitigate batch effects. Fig 5B shows only those individuals where the major cell types were available. The Covid patients originally annotated as Mild/Moderate was included as ‘Moderate’ in Fig 5D.

#### Sepsis-PBMC Dataset (Reyes et al.)

The pre-processed scRNA-seq data was downloaded through the <u>Broad Institute Single Cell Portal</u> <u>(SCP548)</u>. The data was further analyzed with Seurat to obtain the Module score based on UCell.

#### Covid – BALF Dataset (Liao et al.)

The data was accessed from <u>GSE145926</u>. The filtered cell matrix was processed with Seurat with the code as provided by the authors of the original article.

### In vivo Mouse model experiments

For induction of pneumonia, E. coli (DH5α strain), OVA-coated E. coli, grown for 18 h in Luria broth medium at 37 °C, were washed twice (1,000g, 10 min, 37 °C), diluted in sterile isotonic saline and calibrated by nephelometry. Bacteria (75 μl, OD600 = 0.6–0.7) were injected i.t. in anesthetized mice to induce nonlethal acute pneumonia. Infected mice were intravenously (i.v.) injected with 10 μg of CD45-PE on days 0, 1, 3 and 5 to evaluate T cell trafficking towards the lung. Five minutes before sample collection on each of these days, 10 μg of CD45-BV480 was i.v. injected to evaluate blood contamination during lung excision. To specifically assess the expression of membrane markers CD3, CD4, CD8 CXCR4 and CD69 conjugated monoclonal antibodies were used on cell suspensions: CD3-bv711 (145-2C11, 7311597, BD Biosciences, 1:200 dilution); CD4-buv395 (GK1.5, 1097734, BD Biosciences, 1:200 dilution); CD8-AlexaFluor700 (RPA-T8, 9025745, BD Biosciences, 1:200 dilution); anti-CD69-APC (H1.2F3, 9204727, BD Biosciences). Two independent experiments with each 3-4 mice were carried out. Anova test were used to evaluate for statistical significance across time points.

### Analysis of publicly available data set of 10 immune mediated diseases

Bulk RNAseq of 28 pure immune cell types from 339 individuals divided into 10 immune-mediated diseases and 92 healthy controls was obtained from the National Bioscience Database Center (NBDC) Human Database with the accession number <u>E-GEAD-397</u>. The dataset was assembled as a large matrix with genes listed in rows and columns are individuals with homogeneous cell types. Functions from the Seurat pipeline were used to compute the module scores and generate the population’s UMAP embedding.

### Disease Classification

Combined diseases vs healthy approach was deployed for each major cell type and by cell subtype at the primary level. The Logistic Regression classifier from the R package ‘caret’ and the repeated cross-validation strategy for model evaluation were used.

In the next phase, for each major cell type, classification was evaluated for Healthy vs One Disease. An identical classification model was built in this phase as well. The coefficient of variation (CV^2^)^61^ method produced an unsupervised set of highly variable genes as a control for the ALARM module genes. The mean vs (variance/means^2^) was modelled with *glmgam.fit* from the *statmod* R package for the variance estimate of every gene. The genes were ranked by the significance of deviation from the fit. The same number of variable genes was then used in the modules.

### Gene association to disease terms

The association between the ALARM genes and immune disease terms was performed using the disgenet2r (v0.99.2) R package ^51^. The ratio of genes associated with immune diseases to the total number of genes within the modules was compared to the corresponding ratio obtained from 1000 randomly selected sets of 61 genes.

### Methodological Clarifications and Editing

Parts of this manuscript were edited and refined with the assistance of ChatGPT, the AI language model developed by OpenAI. This tool was used to improve the clarity and coherence of the text without altering the scientific content.

## Acknowledgements

We are most grateful to the Genomics Core Facility **GenoA**, member of Biogenouest and France Genomique and to the Bioinformatics Core Facility **BiRD**, member of Biogenouest and Institut Français de Bioinformatique (IFB) (ANR-11-INBS-0013) for the use of their resources and their technical support. DS was supported by Progreffe foundation. The project was supported by Region Pays de Loire “Etoile Montantes” grant for JP. We would like to thank the members of the DIVAT consortium for their involvement in the study, the physicians who helped recruit patients, and all patients who participated in this study. We also thank the clinical research associates who participated in the data collection. Data were collected from the French DIVAT multicentric prospective cohort of kidney and/or pancreatic transplant recipients (www.divat.fr, N° CNIL 914184, ClinicalTrials.gov recording: NCT02900040). We also thank the biological resource centre for biobanking (CHU Nantes, Nantes Université, Centre de ressources biologiques (BB-0033-00040), F-44000 Nantes, France). The authors have no conflicting financial interests.

## Contributions

TL, DS & JP analyzed the data and wrote the manuscript. CF and TL performed scRNA-seq experiments. TL and SV, JB and GB performed pig transplantation model experiments. LBD performed cell sorting experiments. CK, RD, MG and SB collected and provided patient data and kidney transplantation PBMC samples. GT & ND performed and analyzed all transwell experiments. AB & AR performed and analyzed mouse pneumonia experiments and critically revised the manuscript. RJ, SB provided samples, contributed to study design, result interpretation and critically revised the manuscript. All authors contributed to the draft of the manuscript. JP conceived the study and oversaw experiments and analyses.

## Data availability

The data underlying Figures 1 and 2 will be openly available in GEO and embargo will be lifted upon acceptance.

---

**Figure S1.**
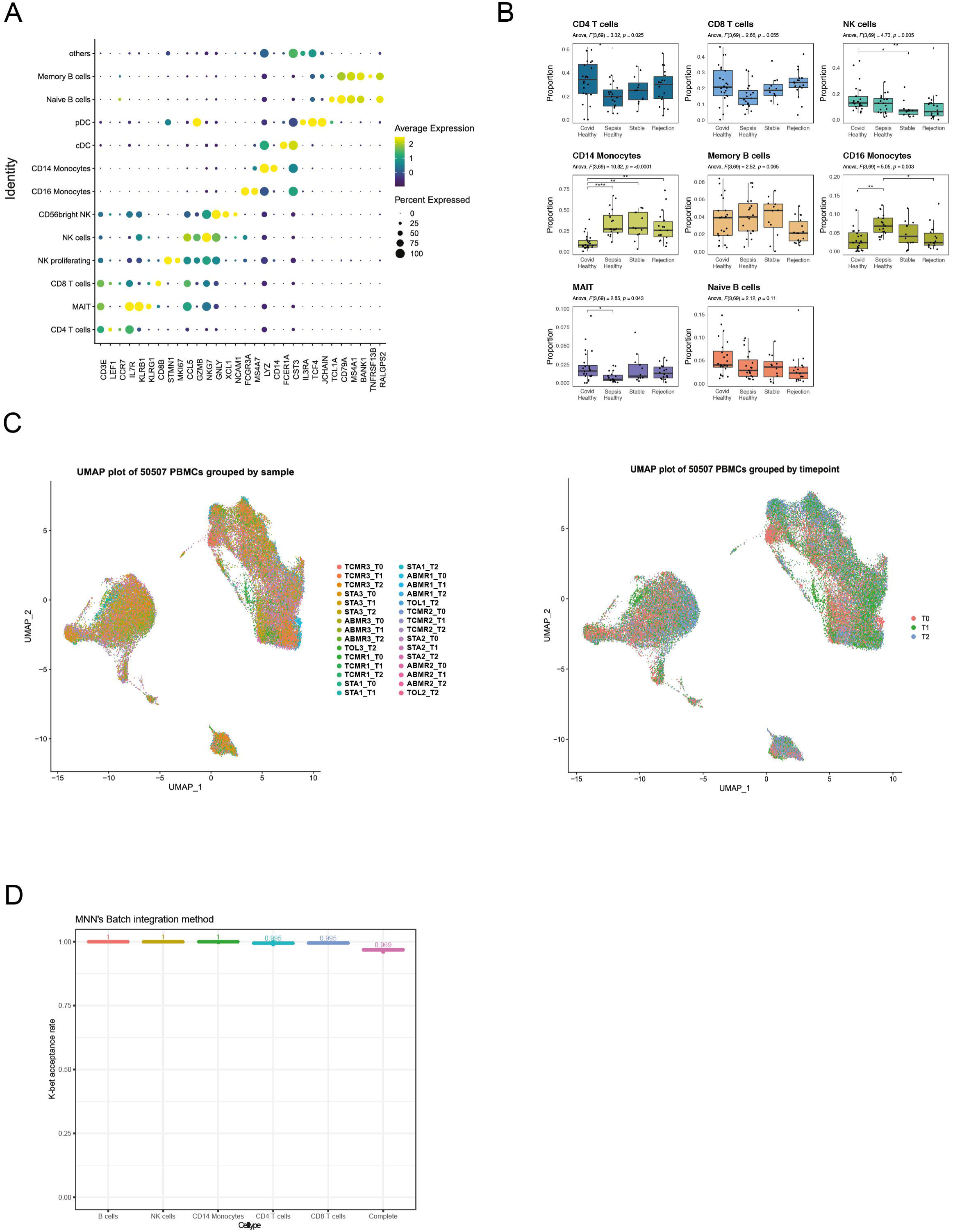
ScRNA-seq integration of a cohort of kidney allograft recipients. A) Expression profiles of cell-specific markers distinguish the PBMC populations. Average expression is the log-normalized expression average of the cells by cell type, size of the dots is associated to the fraction of cells of the cluster in which the gene is detected. B) Cell type proportion by cluster in the overall PBMC population. Comparison include Stable patients, Rejection (ABMR+TCMR) and two public datasets: healthy volunteers from Stephenson et al, 2021, and healthy volunteers from Reyes et al, 2020. One-way ANOVA with Tukey’s multiple comparisons post hoc test was performed, *P < 0.05, **P < 0.01, ***P < 0.001, ****P < 0.0001. C) UMAP projection showing the sample distribution, ABMR=Humoral rejection, TCMR=Cellular rejection, STA=Stable, TOL=Tolerant. Second UMAP shows the timepoint distribution, T0=Time point 0 (Graft), T1=Time point 1, T2=Time point 2. D) K-bet acceptance rate by cell types following the CCA batch integration. Complete k-bet acceptance rate was computed on the overall PBMC population.

**Figure S2.**
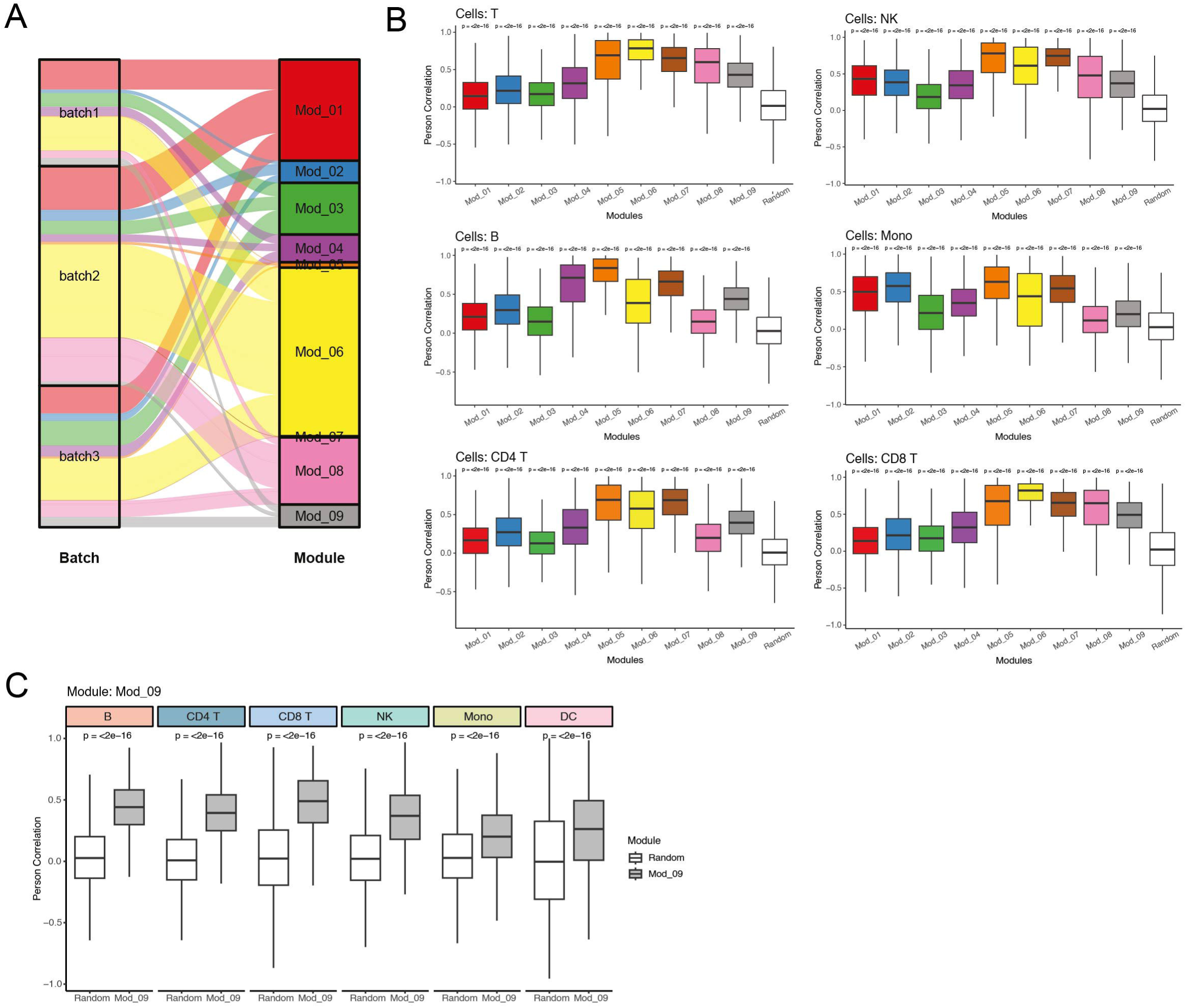
Assessing module distribution. A) Alluvial plot showing the shared origin of each module across batches. Each cell is assigned to the most enriched module they express. B) Boxplots showing the distribution of Pearson correlation coefficient *R* of pairwise gene expression correlation by cell type across modules. A module of random genes was generated to compare with the nine other modules. Multiple gene pairs from each module are selected to compute the rho values using a subset of common cells within the cell type. 1000 repetitions of 50 pairs each yielded 50000 *R* values, as represented in every box. The subset of common cells changes with every repetition. C) Boxplots showing the distribution of Pearson correlation coefficient *R* of pairwise gene expression correlation by cell type for Module 9.

**Figure S3.**
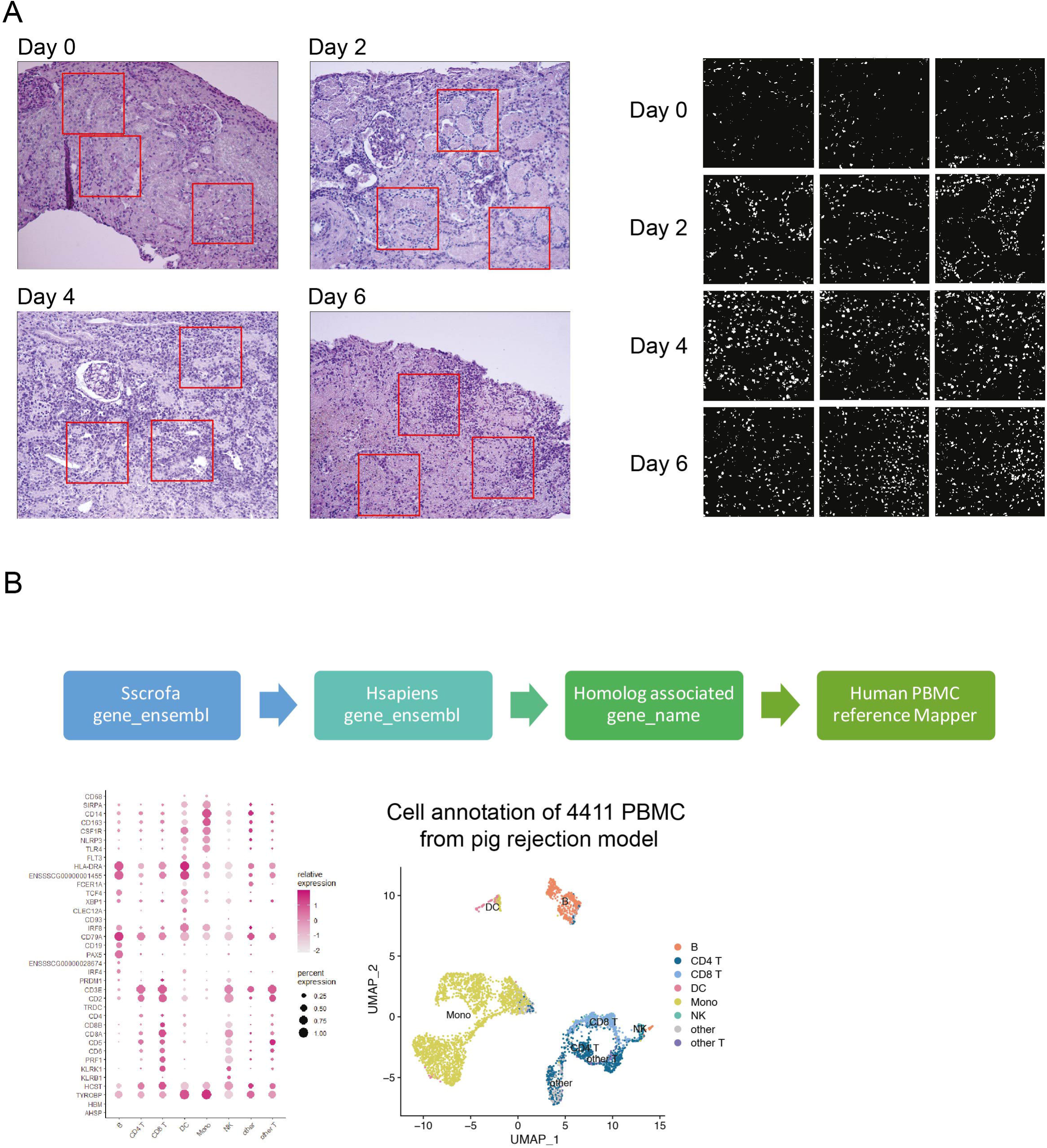
Cellular characterization of pig PBMC by scRNA-seq analysis. A) Image J analysis of immuno-stained slices to quantify cell infiltration. Red boxes show counting areas which excludes glomeruli. Right panel shows the cells which were then counted using Image J software. B) Pig data were annotated using the Sscrofa reference genome and associated to their human homologue. The expression profiles of cell type-specific markers is shown in the dotplot of relative expression by cell type. Size of the dots are associated to the fraction of cells of the cluster in which the gene is detected. The UMAP embeds the 4 samples (D0, D2, D4, D6) with their corresponding cell type annotation.

**Fig S4.**
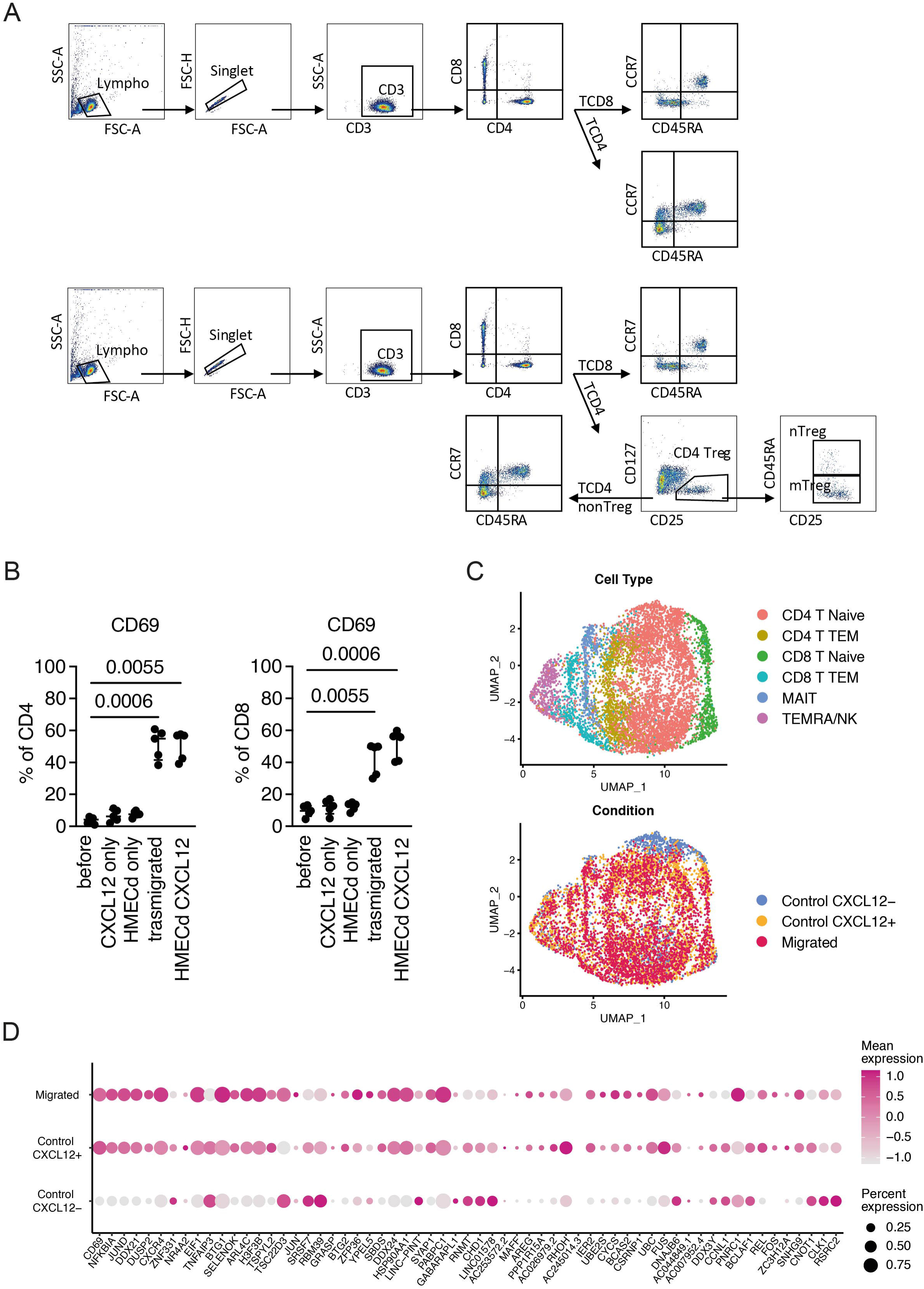
Transwell assay experiments A) Gating strategy used in the transwell assay for cell type annotation. Lymphocytes were gated based on forward scatter (FSC) and side scatter (SSC), followed by singlet selection and the identification of CD3+ T cells. CD4+ and CD8+ T cells were further delineated based on CD45RA and CCR7 expression to distinguish between naïve (Tn), central memory (Tcm), and effector memory RA (TemRA) subsets. Additionally, CD4 T regulatory (Treg) cells were identified by CD25 and CD127 expression to differentiate between activated (aTreg) and non-Treg populations. B) Percentage of CD4+ and CD8+ T cells expressing CD69 under five different experimental conditions: before the transwell assay (baseline), CXCL12 only, HDMEC alone, CXCL12 with HDMEC, and after migration through HDMEC with CXCL12. C) UMAP visualization of scRNA-seq data from the transwell experiment showing all cell populations. The top plot displays T cell subtypes identified by their transcriptional profiles, including CD4+ and CD8+ T naïve cells, CD8+ T effector memory RA (TEMRA) cells, mucosal-associated invariant T (MAIT) cells, and CD4+ and CD8+ T effector memory (TEM) cells annotated using Celltypist. The bottom plot shows the distribution of cells based on their experimental condition: control which is CXCL12-, CXCL12+ and transmigrated. The clustering of cells in the UMAP space indicates differences in transcriptional states based on both cell type and condition. D) ALARM gene expression by condition. Dot plot showing the relative expression levels of ALARM-associated genes in CD4+ and CD8+ T cells across the different conditions of the transwell assay (Migrated, CXCL12+, and CXCL12-). The size of the dots represents the percentage of cells expressing the gene, and the color intensity represents the level of expression, with darker shades indicating higher expression.

**Fig 5S.**
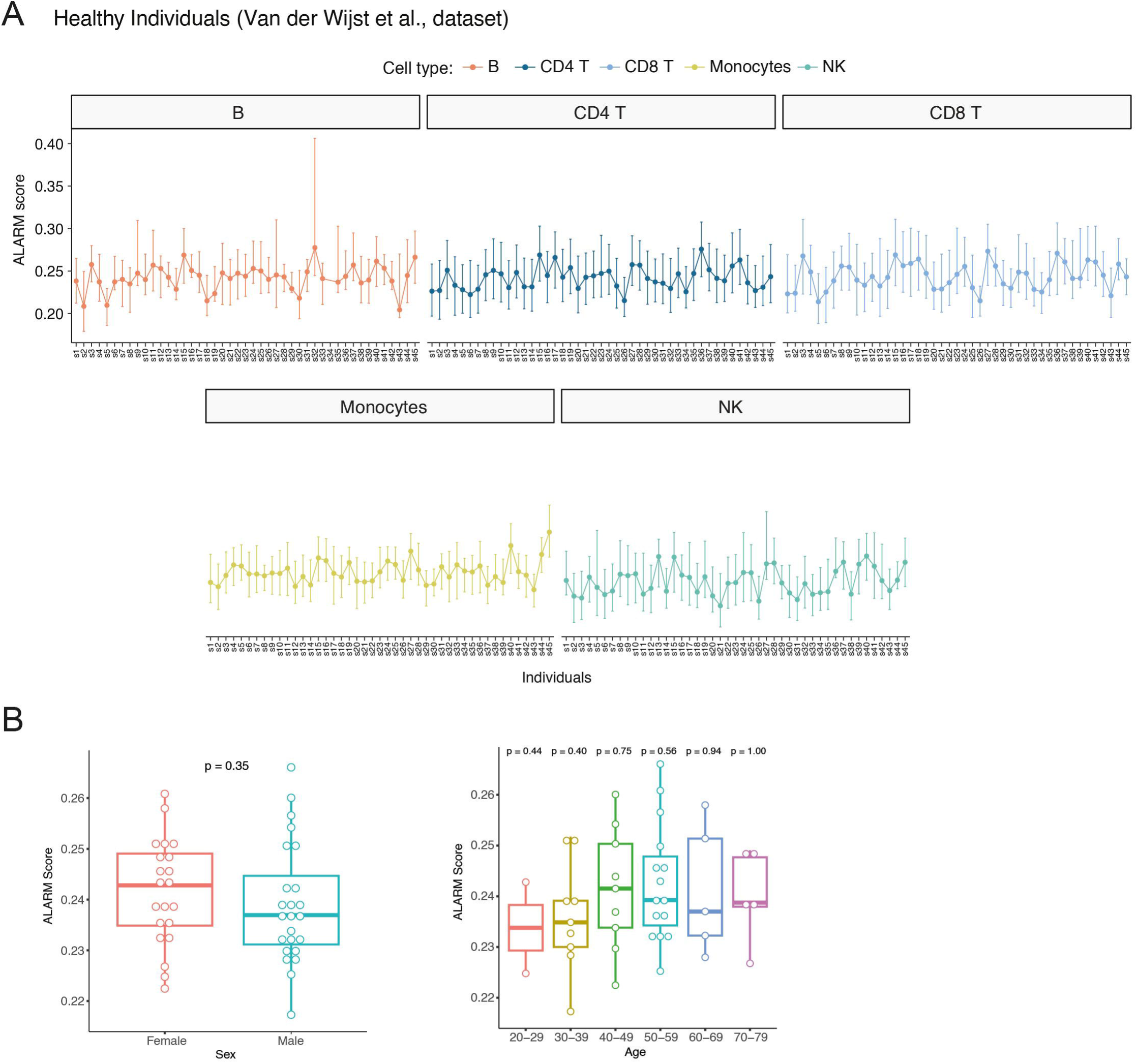
Overview of the cohort distribution A) ALARM module score distribution by cell type based on scRNA-seq analysis of PBMC from 45 unrelated healthy individuals^40^. B) Left: ALARM module score distribution across sex. No significance was found for the sex parameter with a t-test. Right: ALARM module score distribution across age categories. No significance was found for the age parameter with a one-way ANOVA test.

**Fig S6.**
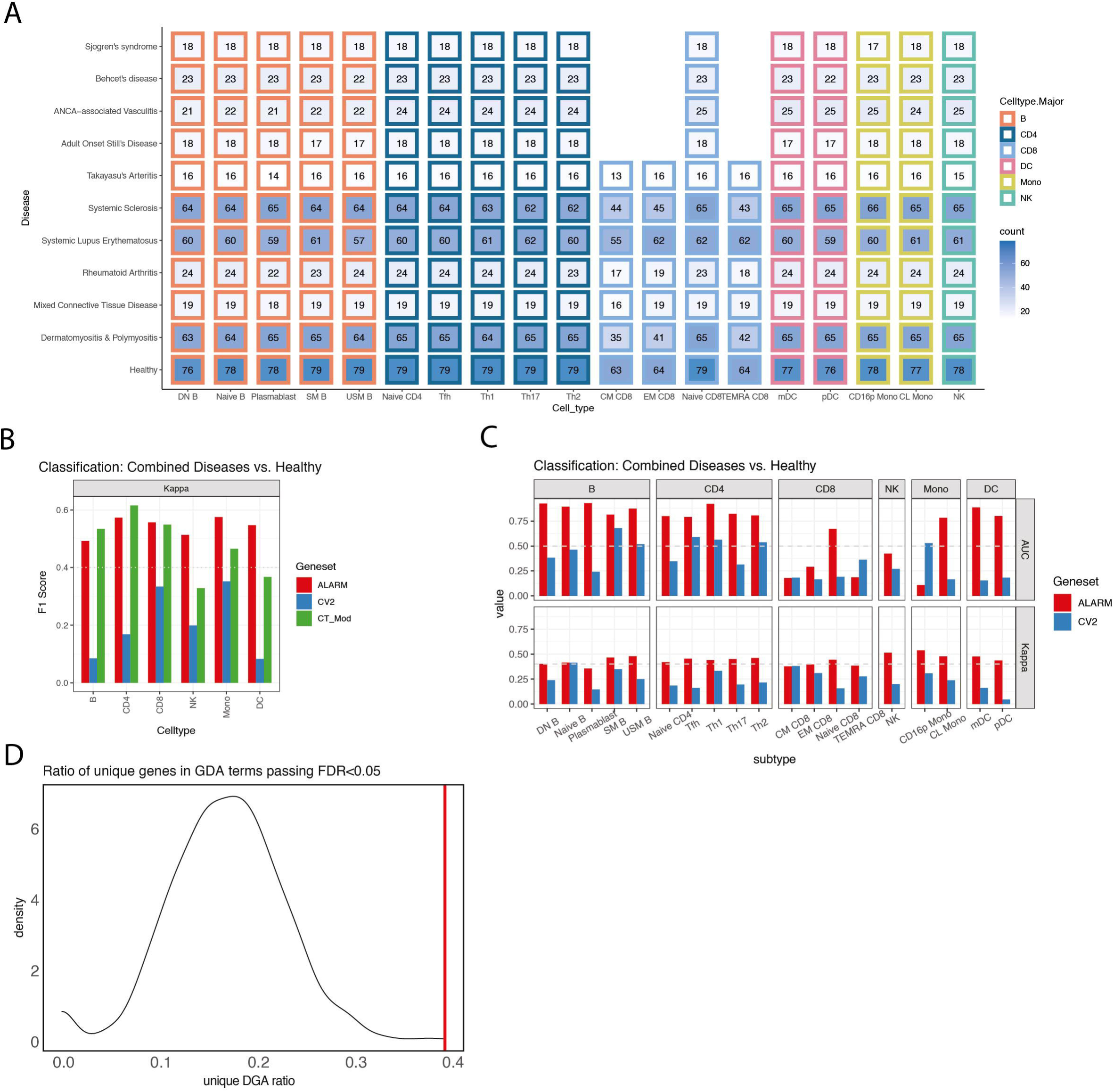
ALARM genes can classify immune-mediated diseases A) Distribution of the bulk RNA-seq samples across the 28 circulating immune cell types for the 337 patients distributed across 10 immune mediated diseases and the 79 healthy controls. B) Area Under Curve (AUC) and Cohen’s Kappa score across cell types in disease and healthy patients. C) AUC and Cohen’s Kappa score across cell subtypes in disease and healthy patients D) Distribution of the ratio of genes associated to Gene-Disease Association (GDA) terms in 1000 modules of 61 randomly selected genes. The ALARM module has a ratio of 0.39 genes associated to GDA terms (red line).

